# Engineered Amphiregulin Promotes Tissue Healing via Dual Regenerative and Immunomodulatory Functions in Tissue-Resident Non-Immune Cells

**DOI:** 10.64898/2026.07.17.738025

**Authors:** Julien M.D. Legrand, Celeste Piotto, Yen-Zhen Lu, Bhavana Nayer, Yuxuan Luo, Sinnee Lau, Jacqueline A. Larouche, Trevor Wilson, Ziad Julier, Mikaël M. Martino

**Affiliations:** Australian Regenerative Medicine Institute, Monash University, Clayton, Victoria 3800, Australia; Biomedical Manufacturing, Commonwealth Scientific and Industrial Research Organisation, Clayton, Victoria 3168, Australia; MHTP Medical Genomics Facility, Monash Health Translation Precinct, Clayton, Victoria 3168, Australia; Victorian Heart Institute, Monash University, Melbourne, Victoria 3800, Australia

## Abstract

Amphiregulin (AREG), a growth factor prominently expressed by immune cells, has emerged as an important mediator of tissue healing. However, its therapeutic potential and immunomodulatory effects remain elusive. Here, we engineered an optimized AREG (eAREG) with enhanced signaling and show that it promotes robust skin repair and muscle regeneration in murine models. Beyond its growth factor function, eAREG acts on tissue-resident non-immune cells to suppress inflammation-induced chemokine programs, thereby limiting the recruitment of pro-inflammatory immune cells. We further demonstrate that eAREG constrains chromatin accessibility at regulatory regions of key chemokine genes, revealing an epigenetic mechanism of immune regulation operating within non-immune tissue compartments. Importantly, the regenerative and immunomodulatory effects of eAREG are preserved in diabetic mice with elevated inflammation and impaired healing. Together, these findings identify eAREG as a dual-function regenerative biologic and establish a design principle for regenerative therapies that integrate morphogenic signaling with immunomodulation mediated by tissue-resident cells.

## Introduction

Successful tissue repair and regeneration following injury depend on the coordinated integration of morphogenic signals and immune regulation (*1-3*). Molecules such as growth factors provide essential morphogenic cues that drive cell proliferation, migration, and differentiation, while immunoregulatory mechanisms, classically attributed to immune cell populations, constrain inflammatory responses to permit progression from injury to repair or regeneration. However, disruption of this balance, particularly through excessive or prolonged inflammation, is a defining feature of impaired tissue healing. For instance, in chronic pathological conditions such as diabetes, tissue healing is frequently compromised due to profound immune dysregulation. Diabetic wounds and muscle injuries are characterized by sustained inflammation, aberrant immune cell recruitment, and defective resolution programs, which collectively impair regeneration and promote fibrosis rather than functional repair (*1-6*). In these settings, the failure to properly integrate morphogenic cues with immune control represents a major barrier to effective tissue healing.

Given their central role in regeneration, growth factors have been extensively explored as therapeutic agents, most commonly delivered as recombinant proteins (*5, 7*). Among these, epidermal growth factor receptor (EGFR) ligands, in particular epidermal growth factor (EGF), continue to be actively investigated for regenerative applications and have been approved in some countries for specific wound healing indications (*8, 9*). However, despite their potency, growth factor therapies often suffer from limited tissue retention, short-lived signaling, potential off-target effects, and insufficient coordination with the immune response, which restricts their efficacy, particularly in inflammatory disease contexts (*5, 7*). Amphiregulin (AREG), another EGFR ligand, has recently emerged as a key mediator of tissue repair and regeneration (*10-16*). In contrast to EGF, AREG is predominantly produced in injured tissues by immune cells, including regulatory T cells (Tregs) and macrophage subsets, possibly positioning it at the interface between growth factor signaling and immune regulation (*10-17*). For example, we and others have shown that AREG expression in Tregs is strongly induced following tissue injury and contributes to repair and regeneration across multiple organs (*10, 12-16*). In addition, we recently identified a unique N-terminal extracellular matrix (ECM)-binding domain in AREG that enables high-affinity binding to matrix components, promoting localized retention and controlled release following delivery (*18*). This ECM-binding mechanism has previously been successfully exploited to engineer growth factors and cytokines with enhanced regenerative efficacy across diverse tissues (*5, 18-20*).

Despite these attractive properties, AREG is known to be a low-affinity EGFR ligand, raising the possibility that its signaling strength may be insufficient to fully harness its regenerative potential when delivered as a recombinant protein. Moreover, the extent to which AREG-driven influences the immune microenvironment during tissue healing, and whether this pathway can simultaneously promote morphogenesis and immune modulation, remains unresolved. Here, we address these questions by first examining the regenerative effect of recombinant AREG and then engineering an optimized form of AREG (eAREG) that enhances EGFR signaling potency while preserving its endogenous ECM-binding capacity. Using models of skin repair and muscle regeneration in both wild-type and diabetic mice, we investigate how eAREG integrates morphogenic signaling with immune regulation to promote tissue healing. Our findings reveal a dual mechanism by which enhanced EGFR signaling in tissue-resident non-immune cells drives regeneration while actively constraining pro-inflammatory immune cell accumulation, establishing a general design principle for regenerative biologics that integrate growth factor potency, spatial persistence, and immune modulation.

## Results

### Recombinant wild-type AREG has limited regenerative effects

To investigate the potential of recombinant AREG to induce tissue healing, two distinct murine models of acute injury were used: full-thickness dorsal splinted excisional wounds for skin repair and volumetric muscle loss for muscle regeneration (*15, 18, 21*), both performed in wild-type mice. For skin wounds, 1 µg of recombinant AREG was delivered by intradermal injection adjacent to the wound on days one and four after injury, resulting in a total dose of 2 µg, which is considered a relatively high dose for growth factor therapies in this model (*19, 20, 22, 23*). Wound closure was assessed at day seven post-wounding, a time point when splinted wild-type wounds remain largely open, unless treated with effective pro-healing interventions (*18*). However, histomorphometric analysis revealed no significant difference in wound closure between the saline control and AREG treatments (**Figure 1A-C**). For muscle injuries, we administered 4 µg of recombinant AREG within a fibrin hydrogel polymerized directly in the defect, a dose that falls within the upper range commonly used for growth factors for this model (*18*). Injuries treated with fibrin hydrogel alone served as the control. The volume of tissue restored was quantified at three weeks post-injury by micro-computed tomography (microCT), and fibrosis was quantified using histology. Similar to the skin model, delivery of recombinant AREG did not enhance muscle restoration compared with control (**Figure 1D-G**). Together, these results show that recombinant AREG does not significantly accelerate tissue healing in skin and muscle injury models, indicating minimal therapeutic potential in this context.

**Figure 1.**
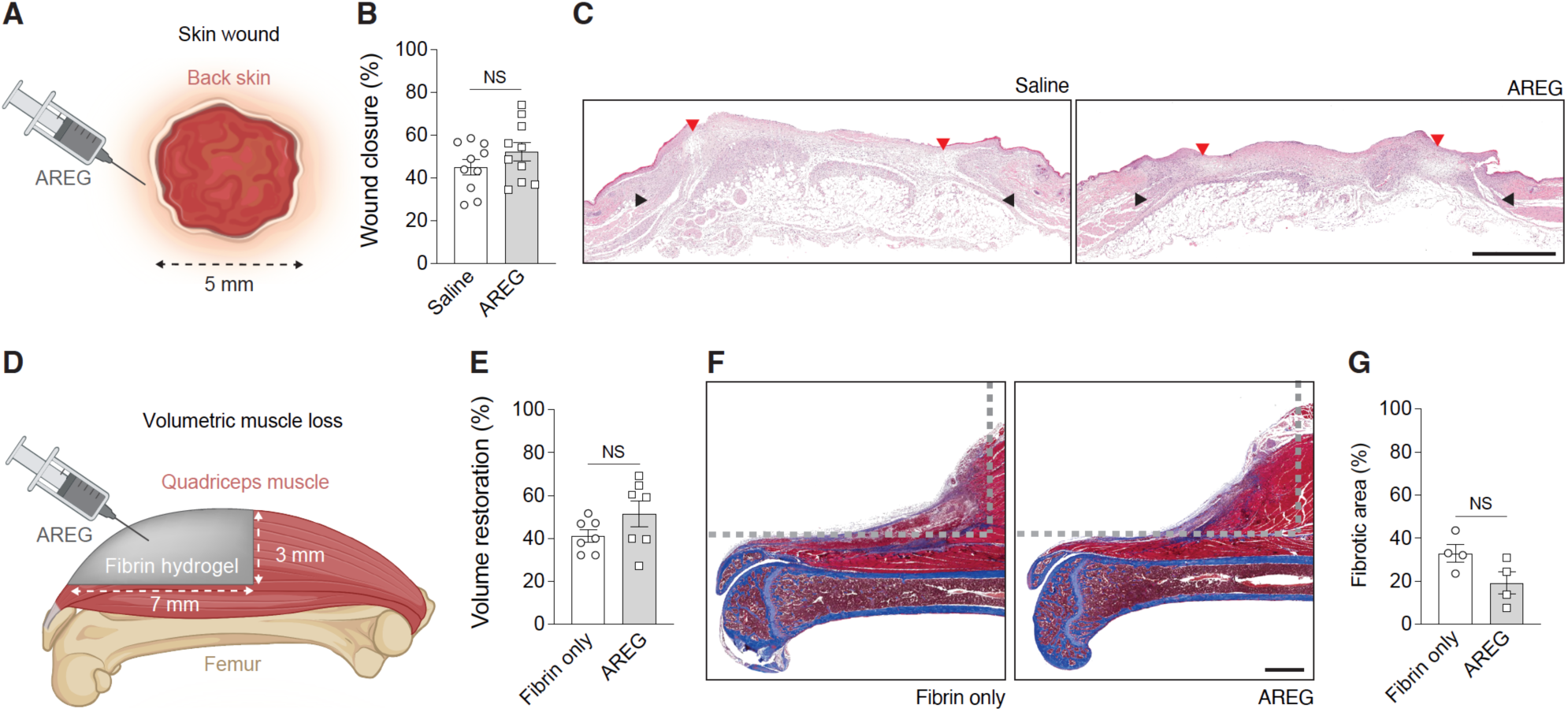
Delivery of recombinant wild-type AREG has limited capacity to promote skin and muscle healing. (**A**) ull-thickness wounds (5 mm diameter) in wild-type mice were treated with intradermal injection of saline or AREG (1 µg total per wound) on days 1 and 4 post-wounding. *n* = 10 wounds. (**B**) Wound closure evaluated by histomorphometric analysis of tissue sections at 7 days post-wounding. (**C**) Representative histology (hematoxylin and eosin staining). Black arrows indicate wound edges, and red arrows indicate tips of epithelium tongue. The epithelium (if any) is stained in purple, underneath which the granulation tissue is stained in pink-violet, with dark purple granulocyte nuclei. Scale bar, 1 mm. (**D**) A volumetric muscle loss defect was created in the quadriceps of wild-type mice and treated with a fibrin hydrogel only or fibrin with AREG (4 µg). (**E**) Tissue volume restoration was measured by microCT 3 weeks post-injury and normalized to the contralateral uninjured muscle on the control leg. *n* = 7. (**F**) Representative histology sections of the center of the muscle defect stained with Masson’s trichrome. Gray dashed lines indicate the original defect. Muscle tissue is stained in dark red, fibrotic parts are in blue/violet, and the femur stained in blue appears under the muscle. Scale bar, 1 mm. (**G**) Quantification of fibrotic tissue in histology sections. *n* = 4. Data are presented as mean ± SEM. Statistical significance was determined using two-tailed unpaired *t* test. NS, not significant.

### Engineered AREG exhibits higher affinity for EGFR and maintains robust ECM-binding capacity

Despite the insignificant regenerative effects observed with recombinant AREG, EGF receptor ligands, including AREG itself, have been shown to support tissue healing in various contexts (*10, 13-16, 24*), suggesting that enhancing AREG activity could yield greater regenerative benefit. Indeed, AREG is known to bind EGFR with relatively low affinity (*25*), which limits the strength of the signaling it can induce. In contrast, high-affinity EGFR ligands such as EGF generate stronger and distinct signaling kinetics (*25, 26*). In addition, we have previously shown that AREG possesses a high-affinity ECM-binding region at its N-terminus that can be isolated and used to improve local retention and regenerative efficacy of various therapeutic proteins (*15*). This ECM-binding property is a major advantage for protein therapeutics delivered directly into injured tissues (*5, 19, 20, 22, 27, 28*). Therefore, we hypothesized that increasing the receptor-binding affinity of AREG, while preserving its endogenous ECM-binding region, would enhance its signaling capacity and thus improve its regenerative potential.

To enhance the interaction of AREG with EGFR while preserving its N-terminal ECM-binding domain, we generated eAREG by partially replacing the C-terminal portion of the AREG receptor-binding domain with the corresponding segment of EGF, the prototypical high-affinity EGFR ligand. This EGF-derived segment contains key residues that confer high-affinity binding to EGFR (*29-31*) and forms part of the conserved EGF-like cysteine-loop domain shared across EGFR ligands, which comprises six cysteine residues (C1–C6) that assemble into three disulfide-stabilized loops essential for receptor engagement (*9*). To maintain the N-terminal ECM-binding domain intact, we selected C2 (Cys147) of the AREG sequence as the substitution point. Amino acids from Ile148 to the C-terminus of mouse AREG were replaced with the corresponding C-terminal amino acids of EGF from Leu991 to Arg1029 (**Figure 2A and Supplementary Figure 1A**). Sequence alignment shows that the engineered construct retained more than 75% homology with mature wild-type AREG (**Supplementary Figure 1B**) and the engineered protein was successfully expressed and purified (**Supplementary Figure 1C**).

**Figure 2.**
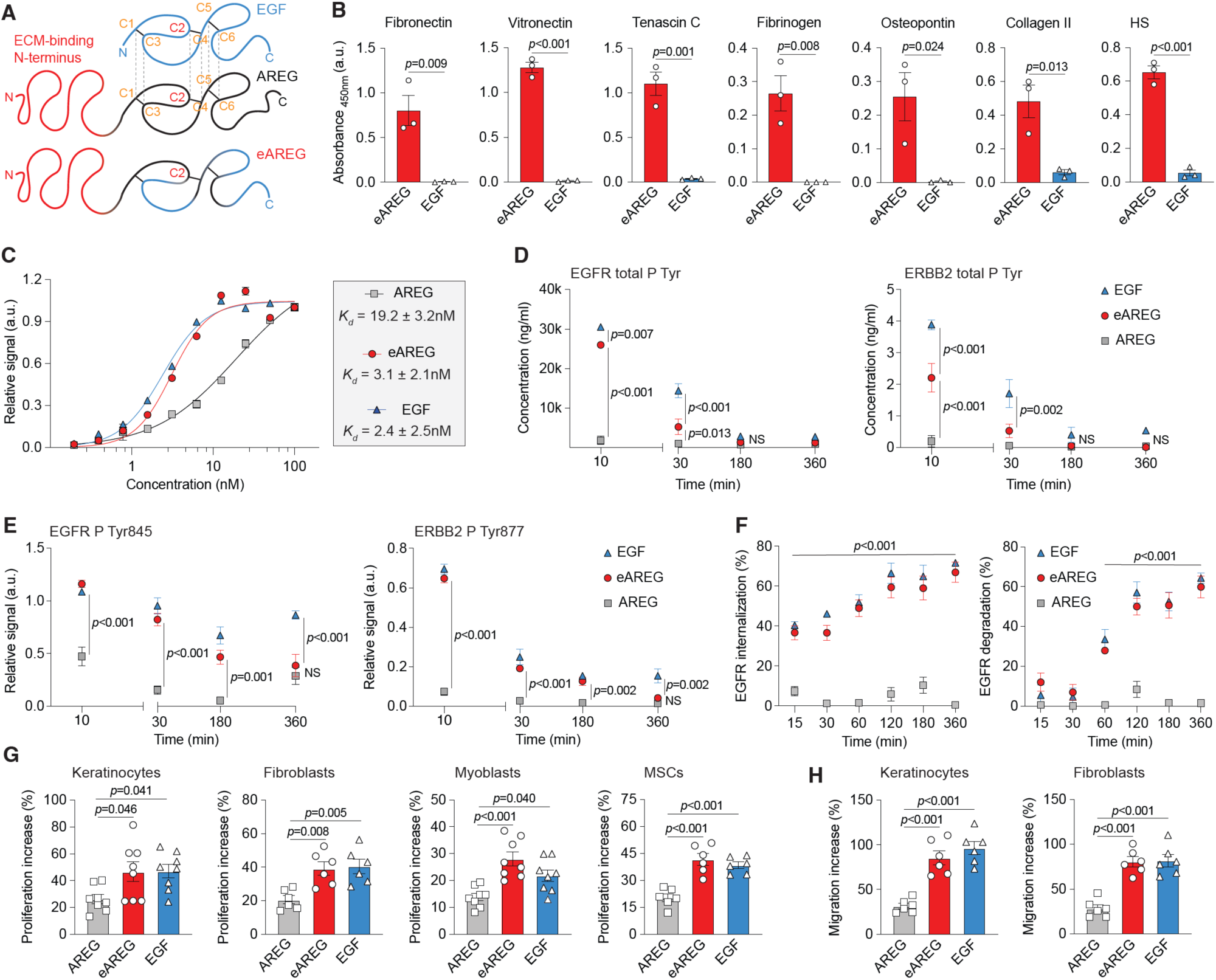
Enhanced signaling by eAREG elicits increased proliferation and migratory responses in tissue cells. (**A**) Engineering strategy to create eAREG. EGF-like domain in EGF, AREG, and eAREG in blue, and blue/black, respectively. ECM-binding domain of AREG in red. N and C indicate the N- and C-terminus, respectively. C1-6 indicate the six cysteines present in the EGF-like domain, creating three disulfide bridges. The black dotted lines connect the equivalent cysteines in the EGF-like domains. The EGF-like domain of AREG is partially substituted with the EGF-like domain of EGF, with C2 (in red) chosen as point of substitution to create eAREG. (**B**) Binding of eAREG and EGF to ECM components. ELISA plates were coated with ECM proteins or heparan sulfate (HS) and incubated with eAREG or EGF. Bound eAREG and EGF were detected with an antibody recognizing the EGF-like domain. *n* = 3 independent experiments. (**C**) Binding affinity of the growth factors to EGFR. ELISA plates were coated with EGFR and incubated with OLLAS-tagged AREG, EGF, or eAREG at increasing concentrations. Bound growth factors were detected with anti-OLLAS antibody. Graphs show the binding curves and the *Kd* obtained from nonlinear regression. *n* = 3 independent experiments. (**D**) Total tyrosine phosphorylation of EGFR and ERBb2 following keratinocyte stimulation with growth factor (100 ng/ml equimolar to EGF) was measured by ELISA after 10, 30, 180, or 360 minutes. *n* = 3 independent experiments. (**E**) Phosphorylation of tyrosine 845 (EGFR) and tyrosine 877 (ERBB2) was measured using an antibody array following keratinocyte stimulation as described in (D). *n* = 3 independent experiments. **(F)** EGFR internalization and degradation was measured by flow cytometry following keratinocyte stimulation with AREG, EGF, or eAREG (100 ng/ml equimolar to EGF) for 15, 30, 60, 120, 180, or 360 minutes. *n* = 3 independent experiments. (**G**) Cell proliferation increase over baseline after 48 hours incubation with AREG, EGF, or eAREG (20 ng/ml equimolar to EGF). *n* = 8 independent experiments for keratinocytes and myoblasts; *n* = 6 independent experiments for fibroblasts and MSCs. (**H**) Transwell cell migration increase over baseline in response to AREG, EGF, or eAREG (20 ng/ml equimolar to EGF) in the bottom chamber. *n* = 6 independent experiments. Data are presented as mean ± SEM. Statistical significance was determined using two-tailed unpaired Student’s *t* test (B), two-way ANOVA with Bonferroni post hoc test for multiple comparisons (D, E, F), and one-way ANOVA with Bonferroni post hoc test for multiple comparisons (G, H). *P* values are indicated. NS, not significant.

To verify that eAREG retained the ability to bind the ECM, the binding of eAREG and EGF to the common ECM components fibronectin, vitronectin, tenascin C, fibrinogen, osteopontin, collagen II, and heparan sulfate was compared using ELISA. eAREG demonstrated strong binding to all ECM proteins tested, including heparan sulfate, whereas EGF, which lacks an ECM-binding domain, showed minimal binding (**Figure 2B**). We also compared eAREG with heparin-binding EGF-like growth factor (HB-EGF), which represents the typical ECM-binding EGFR ligand. eAREG showed markedly stronger binding to ECM components than HB-EGF (**Supplementary Figure 2**). Next, the affinity of eAREG for EGFR compared with EGF and AREG was evaluated by ELISA as an initial validation of the engineering strategy. As expected, the dissociation constant (*K_d_*) of EGF for EGFR was in the nanomolar range and significantly lower than that of AREG. Notably, the *K_d_* of eAREG did not differ significantly from that of EGF, indicating that both ligands bind EGFR with similarly high affinity (**Figure 2C**). Together, these results indicate that eAREG retains strong ECM-binding properties while displaying an increased affinity for EGFR that is comparable to EGF.

### eAREG has EGF-like signaling and greater activity on tissue cells

Because eAREG showed stronger binding to EGFR than AREG, we next investigated its ability to induce phosphorylation of EGFR family receptors. The EGFR family consists of four erythroblastic oncogene B (ERBB) receptor tyrosine kinases: ERBB1 (EGFR), ERBB2, ERBB3, and ERBB4 (*32*), which undergo dimerization and tyrosine phosphorylation upon ligand binding. Although AREG and EGF are ligands specific for EGFR, they can also activate other ERBB receptors through heterodimerization (*25, 33, 34*). Notably, EGF has been reported to preferentially induce EGFR–ERBB2 heterodimers (*35*). Based on this, we assessed total tyrosine phosphorylation (pTyr) of EGFR and ERBB2 as readouts of receptor activation following growth factor stimulation. Keratinocytes, a well-established cellular model for EGFR signaling, were treated with equimolar concentrations of AREG, EGF, or eAREG, and pTyr levels were quantified by ELISA at time points up to six hours post-treatment. At 10 and 30 minutes post-treatment, total pTyr levels were highest in EGF-treated cells for both EGFR and ERBB2, whereas AREG induced minimal phosphorylation at all measured time points. In contrast, eAREG induced significantly higher levels of pTyr for both receptors compared with AREG at 10 and 30 minutes, although these levels remained slightly lower than those induced by EGF (**Figure 2D**). We next examined phosphorylation of specific tyrosine residues across all four ERBB receptors following growth factor treatment using antibody arrays. Consistent with the total pTyr analysis, phosphorylation of specific tyrosine residues in EGFR, ERBB2, and ERBB4 was significantly higher following eAREG treatment than AREG treatment and was comparable to EGF, persisting for up to three hours post-treatment (**Figure 2E and Supplementary Figure 3**). Phosphorylation of ERBB3 was not detected under any condition.

Then, we examined the capacity of eAREG to induce EGFR internalization and degradation, as ligand-dependent differences in receptor trafficking are known to shape the strength and duration of EGFR signaling (*36-39*). Keratinocytes were stimulated with equimolar concentrations of AREG, EGF, or eAREG, and cell surface and intracellular EGFR levels were measured over time using flow cytometry. EGF treatment resulted in approximately 40% EGFR internalization within 15 minutes, increasing to approximately 70% by six hours. In addition, EGF induced approximately 30% EGFR degradation at one hour post-treatment, which increased to approximately 60% by two hours. In contrast, AREG treatment led to significantly lower levels of EGFR internalization and degradation, with maximal internalization and degradation reaching only approximately 10% at three hours and two hours post-treatment, respectively. Notably, eAREG induced EGFR internalization and degradation kinetics that were not significantly different from those induced by EGF at any time point. These findings indicate that eAREG elicits receptor trafficking dynamics characteristic of EGF (**Figure 2F**).

To determine whether differences in EGFR activity induced by eAREG versus AREG have functional consequences for cell behavior, we performed proliferation and migration assays using cell types that are well-known to express EGFR and represent key tissue-resident populations involved in skin repair and muscle regeneration. These included keratinocytes, dermal fibroblasts, myoblasts, and mesenchymal stromal cells (MSCs). AREG, EGF, and eAREG all promoted proliferation across all tested cell types. However, EGF and eAREG induced significantly higher levels of proliferation than AREG, with no significant differences observed between eAREG and EGF (**Figure 2G**). Similarly, transwell migration assays revealed that all growth factors significantly increased migration of keratinocytes and MSCs compared with vehicle control, with EGF and eAREG promoting significantly greater migration than AREG (**Figure 2H**). Overall, these data indicate that eAREG exhibits an EGF-like signaling profile, driving stronger mitogenic and migratory responses than wild-type AREG in tissue cell types that are directly relevant to injury repair and regeneration.

### eAREG promotes skin repair and muscle regeneration

Given that eAREG displayed enhanced EGFR signaling, we next assessed its capacity to promote tissue healing using the same models of skin and muscle injury initially used to evaluate recombinant AREG. Identical treatment regimens were employed, comparing eAREG with saline control and recombinant EGF. In the skin wound model, treatment with eAREG resulted in significantly greater wound closure than either saline control or EGF treatment. Notably, eAREG-treated wounds reached an average closure of approximately 70% by day seven, whereas saline- and EGF-treated wounds achieved approximately 45% and 55% closure, respectively (**Figure 3A,B**). In the muscle injury model, eAREG treatment similarly resulted in significantly enhanced regeneration compared with both saline and EGF groups, with no significant difference observed between saline- and EGF-treated injuries. Nearly twice the volume of tissue was restored in eAREG-treated injuries compared with saline controls, with approximately 80% versus 40% restoration, respectively, while EGF-treated injuries showed approximately 55% restoration (**Figure 3C**). Significantly less fibrosis was observed in both eAREG- and EGF-treated injuries compared with controls, with eAREG treatment resulting in significantly lower levels of fibrosis than EGF (**Figure 3D-E**).

**Figure 3.**
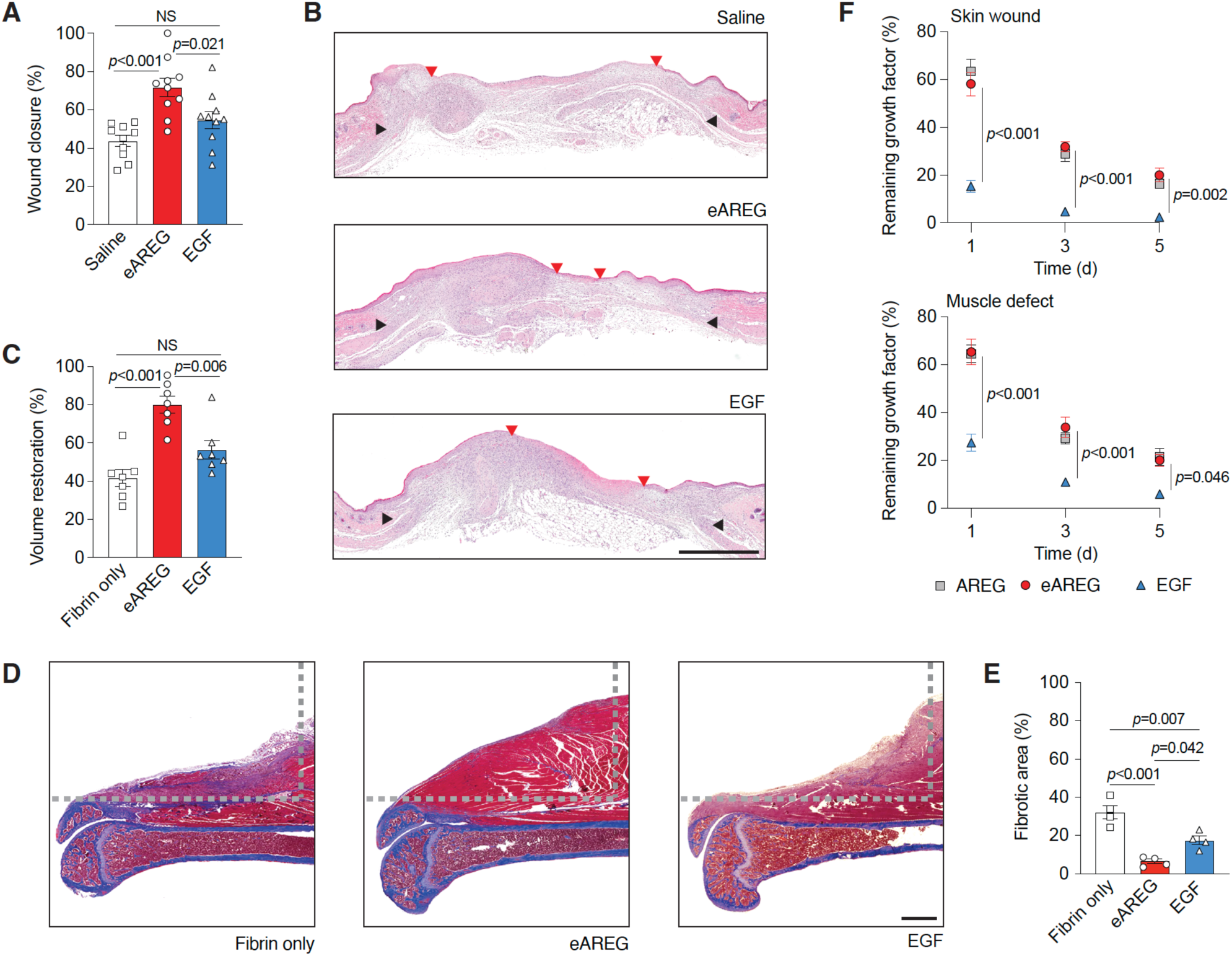
Delivery of eAREG promotes skin repair and muscle regeneration. (**A,B**) Full-thickness dorsal skin wounds (5 mm diameter) were generated in wild-type mice and treated by intradermal injection of saline, eAREG, or EGF (1 µg total per wound, equimolar to AREG) on days 1 and 4 post-wounding. *n* = 10 wounds per group. In A, wound closure quantified by histomorphometric analysis of tissue sections at day 7 post-wounding. In B, representative histological sections stained with hematoxylin and eosin. Black arrows indicate wound edges, and red arrows indicate the tips of the epithelial tongues. The epithelium is stained purple, underlying granulation tissue appears pink-violet, and granulocyte nuclei are dark purple. Scale bar, 1 mm. (**C-E**) A volumetric muscle loss defect was created in the quadriceps of wild-type mice and treated with fibrin hydrogel alone or fibrin containing eAREG or EGF (4 µg, equimolar to AREG). In C, tissue volume restoration measured by micro-computed tomography at 3 weeks post-injury and normalized to the contralateral uninjured muscle. *n* = 7. In D, representative histological sections from the center of the muscle defect stained with Masson’s trichrome. Gray dashed lines indicate the original defect boundary. Muscle tissue is stained dark red, fibrotic tissue blue-violet, and the femur appears blue beneath the muscle. Scale bar, 1 mm. In E, quantification of fibrotic tissue area from histological sections. *n* = 4. (**F**) Retention of AREG, EGF, or eAREG following delivery into skin and muscle injuries. Growth factors were delivered using the same injury models, and the percentage remaining up to 5 days post-delivery was quantified by ELISA. Data are presented as mean ± SEM. Statistical significance was determined using one-way ANOVA with Bonferroni post hoc test for multiple comparisons (A, C, E) and two-way ANOVA with Bonferroni post hoc test for multiple comparisons (F). *P* values are indicated. NS, not significant.

We have previously demonstrated that growth factors and cytokines engineered to incorporate strong ECM-binding sequences can significantly enhance tissue healing by improving their retention at injury sites following delivery (*18-20, 27*). Given that eAREG retains the naturally occurring ECM-binding N-terminal sequence of AREG, we next compared the retention of eAREG, AREG, and EGF at sites of skin and muscle injury following local delivery. Using the same injury models, eAREG, AREG, or EGF were delivered into skin and muscle defects, and the relative amounts remaining at 1, 3, and 5 days post-delivery were quantified by ELISA. Both eAREG and AREG were retained at significantly higher levels than EGF in both skin and muscle injuries at all time points examined, with no significant differences observed between eAREG and AREG. In both skin wounds and muscle injuries, at least 60% of the delivered eAREG and AREG were retained on day 1 post-delivery, whereas retention of EGF was significantly lower (< 20% in skin and < 30% in muscle). By day 5 post-delivery, approximately 20% of eAREG and AREG remained in both skin and muscle injuries, whereas less than 10% of delivered EGF was detected (**Figure 3G**). Together, these findings indicate that eAREG combines enhanced EGFR signaling with strong ECM-mediated retention at injury sites, resulting in superior therapeutic efficacy in acute skin and muscle injury models compared with both AREG and EGF.

### eAREG attenuates inflammatory immune cell accumulation within injured tissues

The immune response to injury is a key regulator of tissue repair, and modulation of immune cell recruitment or activation can strongly influence healing outcomes (*2, 3*). Because AREG is expressed by immune populations within injured tissues, including macrophage subsets and Tregs (*10-16*), EGFR signaling may link growth factor activity to immune regulation during the healing process. We therefore examined how eAREG delivery alters the immune environment of skin and muscle injuries in wild-type mice. Injured tissues were collected for flow cytometric analysis of immune cell composition during the first week post-injury, encompassing the inflammatory and proliferative phases of healing (**Supplementary Figure 4**)(*40*). In skin wounds, we observed a marked reduction in neutrophil numbers following eAREG treatment. Neutrophils, identified as CD11b⁺ F4/80^-^ Ly6G⁺ cells, were reduced by approximately 50% in eAREG-treated wounds compared with saline controls from day 2 through day 7 post-injury. In addition, monocytes/macrophages (Mo/MΦ), defined as CD11b⁺ Ly6G^-^ F4/80⁺ cells, were significantly reduced following eAREG delivery (**Figure 4**). This reduction was most pronounced at early time points, particularly days 2 and 4 post-injury, suggesting attenuation of early pro-inflammatory immune cell recruitment. Consistent with this interpretation, we also observed a significant reduction in CD11b⁺ Ly6C^hi^ cells, indicative of pro-inflammatory Mo/MΦ, within eAREG-treated wounds at these same time points (**Figure 4**). CD206, a marker of pro-resolving Mo/MΦ phenotypes, did not differ between treatment conditions at any time point, suggesting that eAREG primarily affects immune cell recruitment rather than macrophage polarization toward a pro-resolving status (**Supplementary Figure 5**).

**Figure 4.**
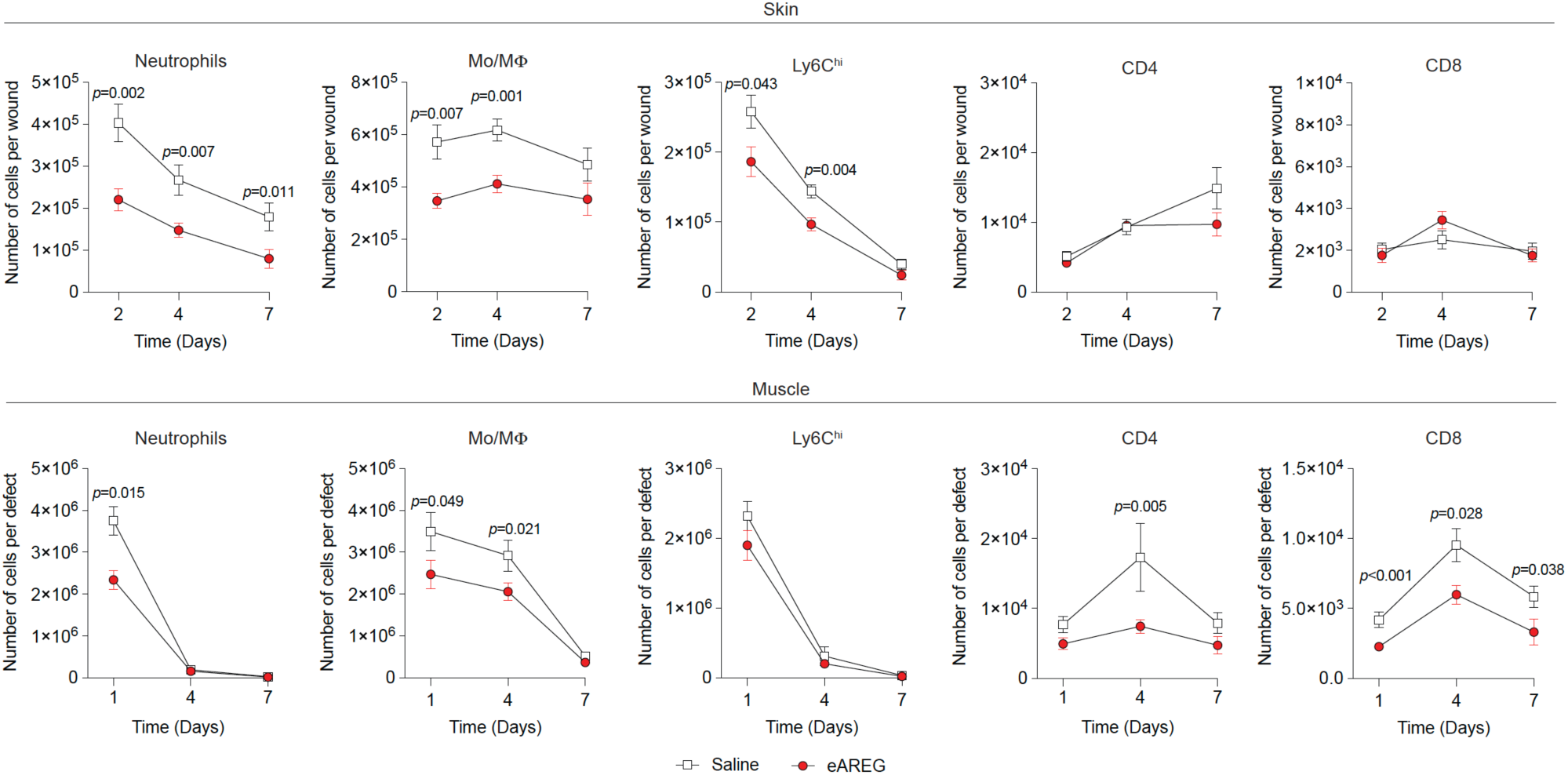
Delivery of eAREG attenuates inflammatory immune cell accumulation within injured tissues. Analysis of neutrophils, Mo/MΦ, pro-inflammatory Mo/MΦ (Ly6C^hi^), CD4^+^ T cells (CD4) and CD8^+^ cytotoxic T cells (CD8) by flow cytometry during tissue healing in wild-type mice. Skin wounds were treated with saline or eAREG on days 1 and 4 post-wounding and analyzed at days 2, 4, and 7 post-injury. Muscle injuries were treated with saline or eAREG and analyzed at days 1, 4, and 7 post-injury. *n* = 10 skin wounds and *n* = 8 muscle injuries. Data are presented as kinetic line plots showing mean ± SEM. Statistical significance was determined using two-way ANOVA with Bonferroni post hoc tests. *P* values are indicated.

A similar immune profile was observed in muscle injuries. eAREG treatment resulted in a significant reduction in neutrophil numbers at day 1 post-injury, as well as reduced Mo/MΦ numbers at days 1 and 4 compared with control treatment (**Figure 4**). As in skin, no differences in Mo/MΦ CD206 expression were detected between conditions (**Supplementary Figure 5**). While no significant changes in CD11b⁺ Ly6C^hi^ cells were observed in muscle injuries, eAREG-treated muscles exhibited a significant reduction in CD4⁺ T cells at day 4 and CD8⁺ cytotoxic T cells at all assessed time points compared with controls (**Figure 4**). In both skin and muscle injury models, no significant differences were observed in the numbers of natural killer (NK) cells, NK T cells, γδ T cells, or B cells between treatment groups (**Supplementary Figure 5**). Collectively, these data indicate that eAREG treatment selectively limits the accumulation of specific pro-inflammatory immune cell populations within injury sites.

### eAREG suppresses chemokine secretion by non-immune tissue cells via epigenetic mechanisms

The reduction in inflammatory immune cell accumulation following eAREG treatment raised the possibility that this effect was mediated by direct signaling to immune cells. However, we found that EGFR expression is very low on major immune cell populations within injured tissues, whereas EGFR is robustly expressed on tissue-resident CD45^-^ non-immune cells (**Supplementary Figure 6A,B**). These findings suggest that the immunomodulatory effects of eAREG are unlikely to result from direct action on immune cells and instead point to an indirect mechanism.

Chemokines are central regulators of immune cell recruitment to sites of injury, where their production by tissue-resident stromal cells following tissue damage promotes immune cell infiltration and subsequent inflammatory responses (*41, 42*). Given the reduced accumulation of pro-inflammatory immune cells observed in eAREG-treated injuries, we investigated whether eAREG modulates inflammation-induced chemokine production by non-immune tissue cells. As an initial screen, keratinocytes, dermal fibroblasts, myoblasts, and MSCs were stimulated with inflammatory cytokines in the presence or absence of eAREG, and chemokine levels in culture supernatants were assessed using membrane-based antibody arrays. Treatment with a cytokine cocktail consisting of tumor necrosis factor alpha (TNF-α), interleukin-1 beta (IL-1β), and interferon gamma (IFN-γ) induced robust chemokine production compared with non-inflammatory controls. Detected chemokines included neutrophil, Mo/MΦ, and T cell chemoattractants such as CCL2, CCL5, CXCL5, CXCL9, CXCL10, and CXCL11 (*43-48*). Notably, co-treatment with eAREG significantly reduced the levels of these chemokines compared with inflammatory cytokine treatment alone (**Supplementary Figure 7**), suggesting that EGFR signaling via eAREG suppresses chemokine production or release. To validate these findings, we performed ELISA to quantify selected chemokines using the same *in vitro* system. Compared with inflammatory cytokine treatment alone, co-treatment with eAREG resulted in significantly reduced levels of CCL2 across all tested cell types; CCL5 in keratinocytes, myoblasts, and MSCs; CXCL9 in keratinocytes, fibroblasts, and myoblasts; and CXCL10 in keratinocytes, fibroblasts, and MSCs (**Figure 5A**), confirming the membrane array results.

**Figure 5.**
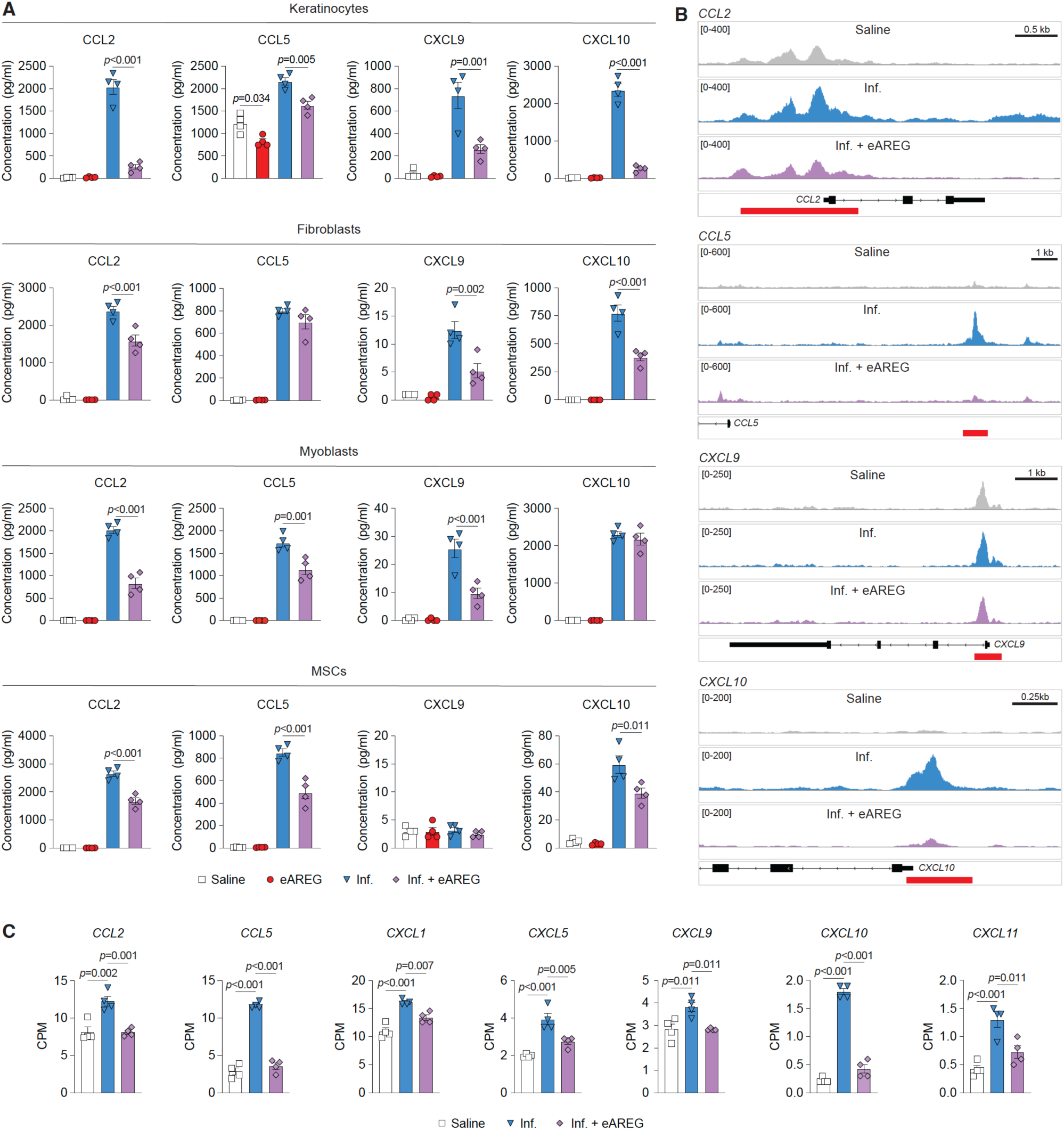
eAREG suppresses chemokine production by non-immune tissue cells and limits chromatin accessibility. (**A**) Keratinocytes, fibroblasts, myoblasts, and MSCs were treated with saline control; eAREG only; an inflammatory cytokine cocktail comprised of TNF-α, IL-1β and IFN-γ only (Inf.); or inflammatory cytokine cocktail and eAREG (Inf. + eAREG). Supernatants were collected after 24 hours and levels of chemokines were quantified by ELISA. *n* = 4 independent experiments. (**B,C**) Keratinocytes were treated with saline control; an inflammatory cytokine cocktail comprised of TNF-α, IL-1β and IFN-γ only (Inf.); or inflammatory cytokine cocktail and eAREG (Inf. + eAREG). Cells were collected after 24 hours and processed for analysis by ATAC-seq. Data were processed using the *nf-core/atacseq* pipeline and regulatory regions (i.e. promoters/enhancers) were annotated using HOMER (as part of the pipeline) or identified within the GeneHancer database. *n* = 4 experimental replicates. In B, merged ATAC-seq read tracks are displayed for each treatment condition at the promoter/enhancer loci for *CCL2*, *CCL5*, *CXCL9* and *CXCL10*. The schematic for each gene is displayed in black below the tracks, and the quantified peaks/regulatory regions are indicated by the red bars. Scale bars are indicated in kilobases (kb). In C, quantification of differences in counts per million mapped reads (CPM) between conditions for ATAC-seq peaks covering annotated regulatory regions (i.e. promoters/enhancers) associated with the displayed chemokine genes. Data are presented as mean ± SEM. Statistical significance was determined using one-way ANOVA with Bonferroni post hoc test for multiple comparisons. *P* values are indicated.

Previous studies have reported that EGFR signaling inhibits pro-inflammatory chemokine expression, likely through epigenetic regulatory mechanisms (*49-53*). We therefore used assay for transposase-accessible chromatin using sequencing (ATAC-seq) to test whether eAREG suppresses chemokine expression by limiting chromatin accessibility within the regulatory regions of chemokine genes. Keratinocytes were used as a cellular model and treated with saline control, the inflammatory cytokine cocktail (TNF-α, IL-1β, and IFN-γ), or inflammatory cytokines with eAREG co-treatment prior to ATAC-seq profiling. Inflammatory cytokine stimulation induced a robust increase in chromatin accessibility, measured as ATAC-seq counts per million (CPM), at promoter and enhancer regions associated with multiple chemokine genes compared with saline-treated controls. These loci included key chemoattractants for neutrophils, monocytes, and T cells, such as *CCL2*, *CCL5*, *CXCL1*, *CXCL5*, *CXCL9*, *CXCL10*, and *CXCL11*. Notably, co-treatment with eAREG completely prevented this inflammation-induced increase in chromatin accessibility at chemokine-associated regulatory regions (**Figure 5B,C and Supplementary Figure 8**). Taken together, these findings demonstrate that, in physiological injury models, eAREG suppresses inflammation-induced chemokine programs and the accumulation of pro-inflammatory immune cells through EGFR signaling in tissue-resident non-immune cells.

### Delivery of eAREG promotes tissue healing and suppresses chemokine-driven immune cell recruitment in diabetic mice

Clinical conditions associated with immune dysfunction are known to impair tissue repair and regeneration (*54*). In particular, diabetes induces a chronic inflammatory state that markedly compromises skin wound healing and muscle regeneration (*3, 4, 6*). Given that eAREG promotes tissue healing and attenuates inflammatory immune cell accumulation in wild-type mice, we next investigated its effects in a disease setting characterized by dysregulated immune responses. To this end, we performed skin wound and volumetric muscle loss injuries in obese *db/db* mice, a clinically relevant model of type 2 diabetes (*55, 56*). For diabetic skin wounds, 2 µg of eAREG or an equivalent volume of saline was delivered by intradermal injection adjacent to each wound on days 1, 4, and 7 post-injury. Histomorphometric analysis at day 10 post-injury revealed significantly greater wound closure in eAREG-treated wounds compared with saline-treated controls. Notably, approximately half of the eAREG-treated wounds were fully closed by this time point, with overall closure exceeding 80% of the original wound area, whereas saline-treated wounds remained less than 60% closed (**Figure 6A-C**). For muscle injuries, 4 µg of eAREG or saline control was delivered via fibrin hydrogel at the time of surgery. At three weeks post-injury, eAREG-treated mice exhibited significantly greater tissue restoration compared with saline-treated controls (approximately 70% versus 40% tissue restoration, respectively) (**Figure 6D,E**). In addition, eAREG treatment resulted in significantly reduced fibrosis within the muscle defect compared with saline-treated controls (**Figure 6F,G**). Collectively, these results show that eAREG retains potent pro-healing activity in the diabetic context.

**Figure 6.**
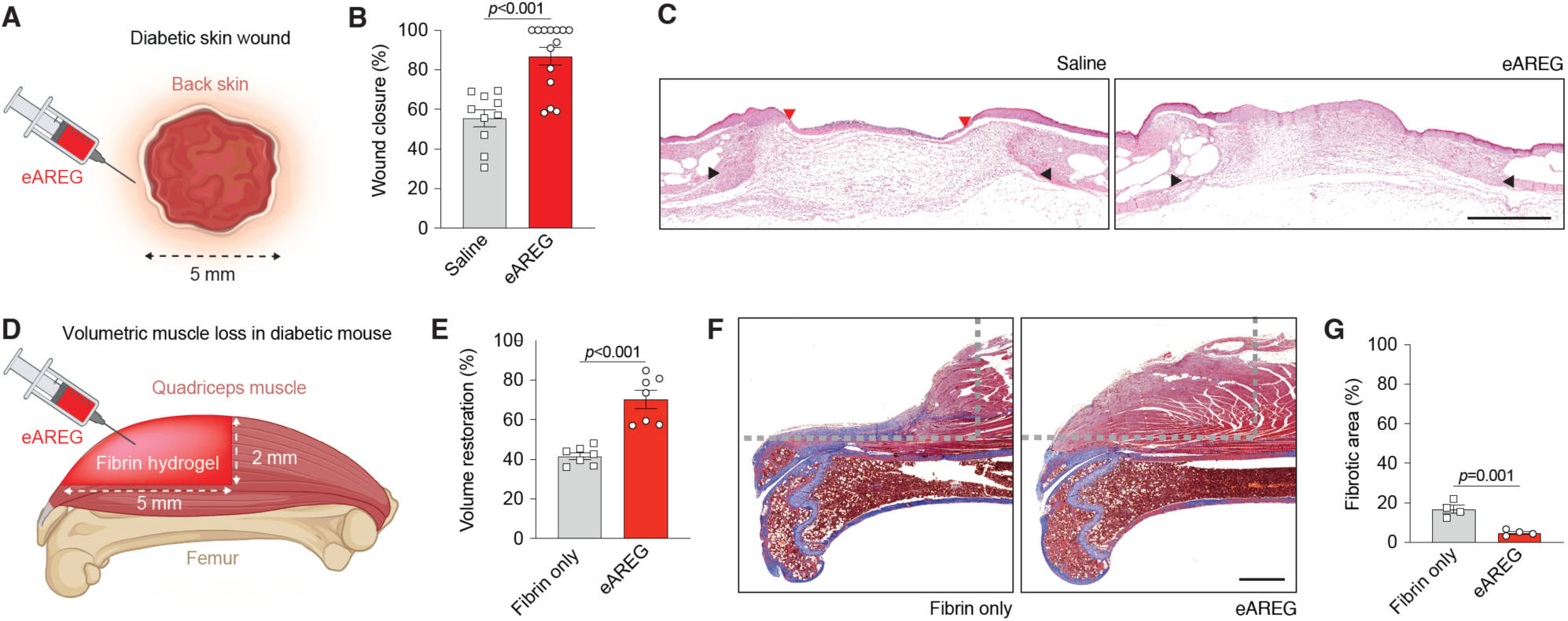
Delivery of eAREG promotes skin repair and muscle regeneration in diabetic mice. (**A**) Full-thickness dorsal skin wounds (5 mm diameter) were generated in *db/db* mice and treated by intradermal injection of saline or eAREG (2 µg total per wound) on days 1, 4, and 7 post-wounding. *n* = 10 wounds for saline and *n* = 14 for eAREG. (**B**) Wound closure quantified by histomorphometric analysis of tissue sections at day 10 post-wounding. (**C**) Representative histological sections stained with hematoxylin and eosin. Black arrows indicate wound edges, and red arrows indicate the tips of the epithelial tongues. The epithelium is stained purple, underlying granulation tissue appears pink-violet, and granulocyte nuclei are dark purple. Scale bar, 1 mm. (**D**) A volumetric muscle loss defect was created in the quadriceps of *db/db* mice and treated with fibrin hydrogel alone or fibrin containing eAREG (4 µg). (**E**) Tissue volume restoration measured by micro-computed tomography at 3 weeks post-injury and normalized to the contralateral uninjured muscle. *n* = 7. (**F**) Representative histological sections from the center of the muscle defect stained with Masson’s trichrome. Gray dashed lines indicate the original defect boundary. Muscle tissue is stained dark red, fibrotic tissue blue-violet, and the femur appears blue beneath the muscle. Scale bar, 1 mm. (**G**) Quantification of fibrotic tissue area from histological sections. *n* = 4. Data are presented as mean ± SEM. Statistical significance was determined using two-tailed unpaired Student’s t test. *P* values are indicated.

We next examined whether the immunomodulatory mechanism of eAREG observed in wild-type mice is preserved in the diabetic model. Chemokine levels were quantified by ELISA in diabetic skin wounds and volumetric muscle loss injuries at day 2 post-injury. Levels of CCL2, CCL5, CXCL9, and CXCL10 were reduced by up to 50% in eAREG-treated skin and muscle injuries compared with controls (**Figure 7A**). Consistent with lower level of chemokines, eAREG-treated skin wounds exhibited reduced numbers of neutrophils, Mo/MΦ, and CD11b⁺ Ly6C^hi^ cells at day 4 post-injury. In diabetic muscle injuries, eAREG treatment similarly resulted in reduced neutrophil numbers at days 2 and 4, as well as decreased Mo/MΦ and CD11b⁺ Ly6C^hi^ cell numbers at day 4 compared with controls. In addition, both CD4⁺ and CD8⁺ T cell numbers were significantly reduced in eAREG-treated muscle injuries at day 2 post-injury (**Figure 7B and Supplementary Figure 4**). These findings demonstrate that the immunomodulatory effects of eAREG are preserved in diabetic injuries and are associated with suppression of chemokine expression and reduced recruitment of pro-inflammatory immune cells.

**Figure 7.**
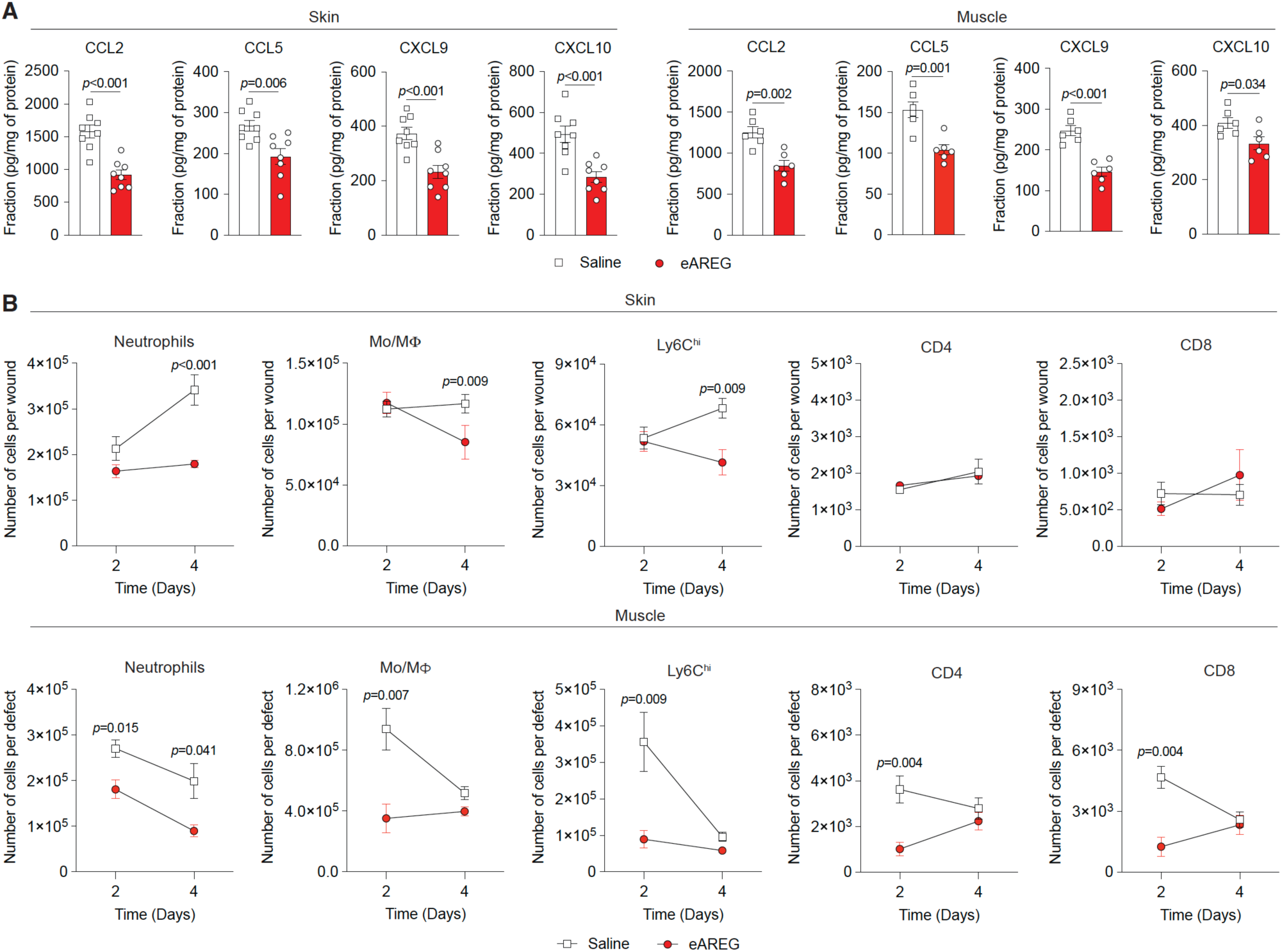
eAREG treatment reduces chemokine production and pro-inflammatory immune cell infiltration in diabetic tissue injury. (**A**) Chemokine levels 2 days after skin and muscle injuries from diabetic mice treated with saline or eAREG were quantified by ELISA. *n* = 8 skin wounds and *n* = 6 muscle injuries. (**B**) Flow cytometric analysis of neutrophils, Mo/MΦ, CD11b⁺ Ly6C^hi^ cells, CD4⁺ T cells, and CD8⁺ T cells in skin and muscle injuries from diabetic mice at days 2 and 4 post-injury. *n* = 10 skin wounds and *n* = 6 muscle injuries per time point. For both panels, skin wounds received saline or eAREG (2 µg) and muscle injuries received saline or eAREG (4 µg). Data are presented as mean ± SEM. Statistical significance was determined using a two-tailed unpaired Student’s *t* test (A) and two-way ANOVA with Bonferroni post hoc tests (B). *P* values are indicated.

Overall, our data support a unified model in which eAREG promotes tissue healing across physiological and disease contexts by enhancing morphogenic signaling in tissue-resident cells and suppressing chemokine-driven pro-inflammatory immune cell recruitment. Importantly, these immunoregulatory effects operate within non-immune tissue compartments rather than via direct modulation of immune cells (**Figure 8**).

**Figure 8.**
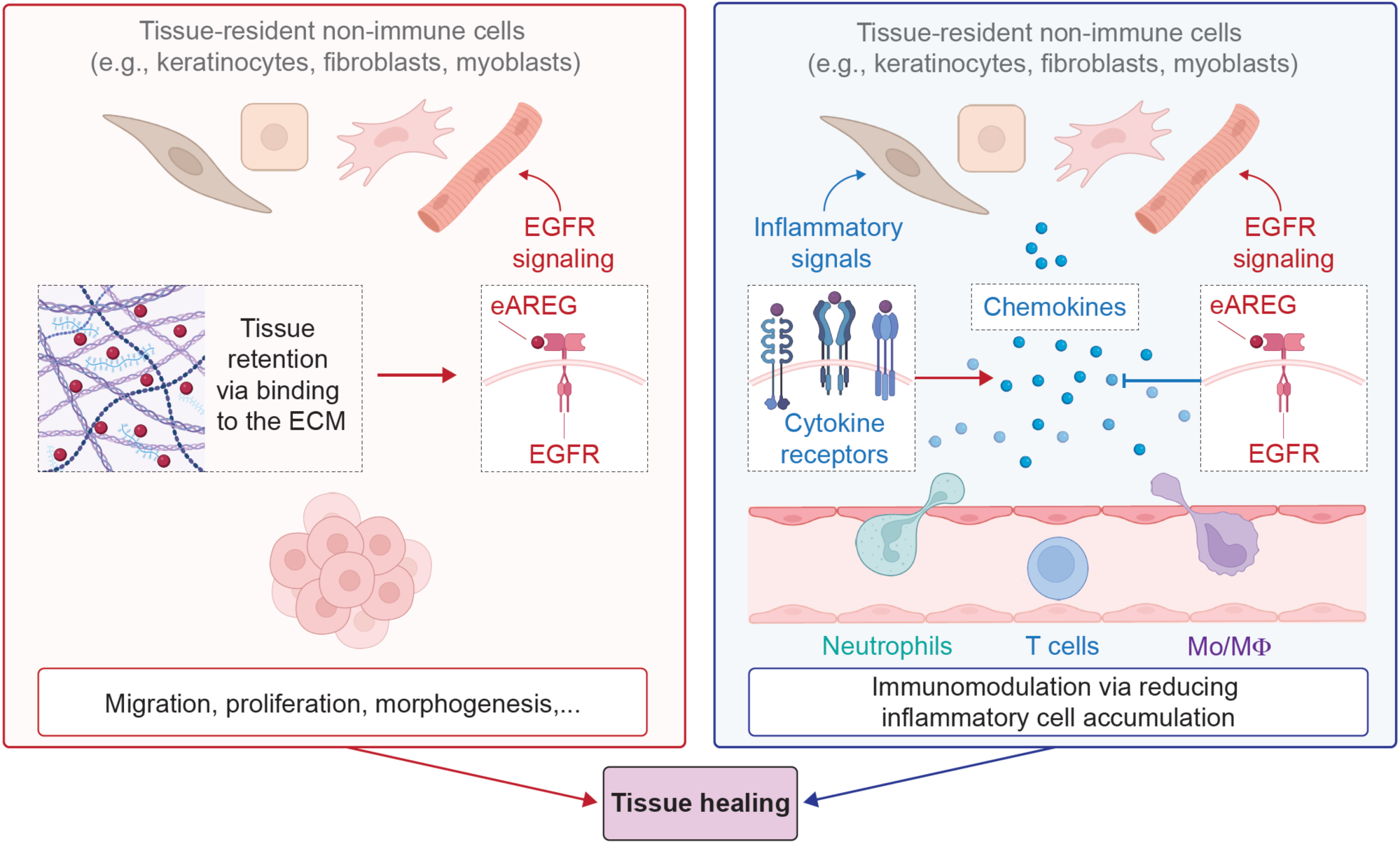
EGFR signaling via eAREG promotes tissue healing via dual morphogenic signals and immunoregulation. EGFR signaling activated by eAREG in tissue-resident non-immune cells stimulates classical morphogenic programs such as cell proliferation and migration, thus directly promoting tissue regeneration. In parallel, EGFR signaling via eAREG suppresses the expression of pro-inflammatory chemokines (e.g., CCL2, CCL5, CXCL9, and CXCL10) that are normally induced by inflammatory cytokines released after tissue injury, including IL-1β, TNF-α, and IFN-γ. This reduction in chemokine production limits the recruitment and accumulation of inflammatory neutrophils, Mo/MΦ, and T cells within injured tissues. Both the pro-morphogenic and immunomodulatory effects of eAREG are enhanced by its strong ECM-binding capacity, which promotes sustained retention at the site of delivery. Together, these mechanisms enable eAREG to promote tissue healing through coordinated regenerative signaling and immunomodulation within tissue-resident cells. Created with BioRender.com.

## Discussion

Morphogenic growth factor signaling and immune regulation are both central determinants of tissue repair and regeneration. While growth factors have long been viewed as the primary drivers of morphogenesis, it is now well established that immune responses critically shape healing outcomes across tissues (*1-3, 57*). In particular, we and others have shown that Tregs play a central role in coordinating tissue repair across multiple organs by providing pro-resolving signals that support effective healing (*10, 12-16*). Notably, AREG is one of the few growth factors robustly expressed by Tregs accumulating in injured tissues, including skin and skeletal muscle (*10, 12, 15*). In addition to this immunological association, AREG possesses a unique N-terminal ECM-binding domain that enables local retention following delivery (*18*).

Although AREG has been implicated in tissue healing across multiple organs, it is a relatively low-affinity ligand for EGFR, raising the possibility that its endogenous signaling properties are insufficient to drive robust regeneration when delivered as a recombinant therapeutic. Consistent with this limitation, wild-type AREG failed to improve healing in both skin wounds and volumetric muscle loss injuries, despite dosing in the standard-to-high range commonly used for growth factors in these models. This apparent disconnect highlights an important distinction between endogenous growth factor function and therapeutic efficacy. Endogenous AREG is produced by Tregs and macrophage subsets within injured tissues and likely operates in highly localized immune-stromal niches, where it is delivered in close spatial and temporal proximity to EGFR-expressing tissue-resident cells. In this context, even low-affinity EGFR engagement may be biologically effective, particularly when combined with sustained local availability and coordinated immune-derived cues. By contrast, recombinant AREG delivered as a soluble protein must rely on receptor binding alone, and its weak receptor phosphorylation and limited receptor trafficking dynamics are likely insufficient to achieve the magnitude and duration of EGFR signaling required to overcome inflammation-induced inhibitory programs in injured tissues. Moreover, EGFR ligands are known to elicit ligand-specific receptor activation and trafficking behaviors (*39*), suggesting that the qualitative features of AREG-induced signaling may be optimized for physiological regulation rather than therapeutic intervention. To overcome these constraints, we engineered AREG to enhance EGFR engagement while preserving its endogenous ECM-binding capacity. By partially substituting the C-terminal portion of the AREG receptor-binding domain with the corresponding region of EGF, we generated eAREG with EGF-like affinity and signaling kinetics. eAREG induced stronger EGFR, ERBB2, and ERBB4 phosphorylation, exhibited receptor internalization and degradation dynamics similar to EGF, and drove robust proliferative and migratory responses in tissue-resident cell types relevant to repair and regeneration.

The enhanced signaling properties of eAREG translated into markedly improved skin repair and muscle regeneration. EGFR signaling in tissue-resident non-immune cells, including keratinocytes, fibroblasts, myoblasts, and MSCs, is classically associated with morphogenic programs such as proliferation, migration, and differentiation. Consistent with this, EGF has been extensively explored in regenerative medicine and is approved for specific topical or local clinical applications in wound healing in selected countries (*8, 58*). Notably, however, eAREG outperformed EGF in both skin and muscle injury models, despite inducing comparable levels of EGFR activation *in vitro*. This superior efficacy is best explained by the strong ECM-binding capacity retained by eAREG, which resulted in prolonged local retention at injury sites compared with EGF. Thus, these findings highlight the importance of integrating signaling potency with spatial persistence to achieve effective tissue repair and regeneration.

Interestingly, we revealed that eAREG treatment had a strong immunoregulatory effect at the injured sites, which led to reduced accumulation of pro-inflammatory immune cells. Indeed, excessive or prolonged infiltration of neutrophils, pro-inflammatory Mo/MΦ, and cytotoxic T cells is well known to impair tissue repair by sustaining inflammatory signaling, promoting ECM degradation and fibrosis, and inhibiting regenerative cell function. For instance, we and others have shown in both skin and skeletal muscle that persistent neutrophil and pro-inflammatory Mo/MΦ activity is linked to delayed healing and defective regeneration, while excessive CD8⁺ T cell accumulation can exacerbate IFN-driven inflammation (*10, 14, 15, 21*). Thus, the ability of eAREG to selectively limit inflammatory immune cell accumulation is likely to create a tissue environment that favors repair and regeneration rather than chronic inflammation. Although we did not test whether wild-type AREG or EGF exert comparable immunomodulatory effects in the injury models, it is likely that both ligands engage similar regulatory pathways downstream of EGFR. However, the relatively weak signaling activity of AREG and the limited tissue retention of EGF likely restrict the magnitude and duration of any such effects *in vivo*. In contrast, the enhanced signaling potency and prolonged tissue persistence achieved with eAREG appear to be required to effectively suppress inflammation-induced chemokine programs and meaningfully alter immune cell recruitment within injured tissues.

Mechanistically, our data indicate that the immunomodulatory effect of eAREG is unlikely to result from direct signaling to immune cells, as EGFR expression was minimal across immune populations within injured tissues. Instead, eAREG acts on tissue-resident non-immune cells to suppress inflammation-induced chemokine production. We found that eAREG reduced the expression and tissue concentrations of key chemokines involved in the recruitment of neutrophils, Mo/MΦ, and T cells, including CCL2, CCL5, CXCL9, CXCL10, and others. This is consistent with previous reports demonstrating that activation of EGFR signaling suppresses pro-inflammatory and IFN-inducible chemokine programs in cancer, whereas EGFR inhibition induces the opposite pattern and is associated with heightened inflammatory chemokine expression and immune cell infiltration (*51*). More broadly, EGFR signaling in epithelial and stromal compartments has been shown to shape the tumor immune microenvironment by regulating chemokine expression and immune cell recruitment, positioning EGFR as an upstream regulator of immune infiltration rather than a direct modulator of immune cell function (*52*). The specific chemokines suppressed by eAREG are well-established drivers of inflammatory immune cell trafficking. CCL2 and CCL5 are known to promote recruitment of Mo/MΦ, whereas CXCL9 and CXCL10 act as IFN-inducible chemoattractant for activated T cells. Importantly, regulation of CXCL9 and CXCL10 downstream of EGFR signaling has been mechanistically linked to epigenetic repression, including EGFR-dependent histone deacetylation at chemokine gene loci, in inflamed epithelial contexts such as human lung adenocarcinoma (*53*).

Our findings directly connect EGFR activation to chromatin-level control of chemokine expression and immune cell dynamics, suggesting that suppression of chemokine networks underlies the reduced accumulation of pro-inflammatory immune cells observed following eAREG treatment. Our ATAC-seq data show that eAREG prevents inflammation-induced increases in chromatin accessibility at promoter and enhancer regions associated with multiple chemokine genes, providing direct evidence that EGFR signaling via eAREG constrains inflammatory chemokine programs at the level of chromatin regulation. While the precise downstream signaling pathways linking EGFR activation to chromatin remodeling were not defined here, EGFR-dependent regulation of epigenetic states has been reported previously and may involve MAPK-, PI3K-, or STAT-dependent signaling pathways that converge on histone-modifying enzymes and transcriptional regulators (*59-61*). It is also plausible that other EGFR ligands, or growth factor families engaging similar signaling architectures, could activate comparable immunoregulatory programs under specific inflammatory conditions. In addition, while this epigenetic mechanism of chemokine suppression was conserved across tissues, its downstream immunological consequences differed in a tissue-dependent manner, with reduced T cell accumulation evident in muscle but not skin, likely reflecting differences in the relative contribution and timing of adaptive immune responses during repair in these distinct tissue contexts.

Interestingly, the immunoregulatory role uncovered here is consistent with the well-documented inflammatory toxicities observed in patients treated with EGFR-targeted cancer therapies. EGFR inhibitors frequently induce inflammatory dermatologic toxicities, including papulopustular rash, xerosis, and barrier dysfunction, in a high proportion of treated patients. These adverse effects are widely attributed to disruption of EGFR signaling in epidermal and stromal cells rather than direct effects on immune cells. Mechanistically, anti-EGFR drugs trigger type I interferon signaling in keratinocytes and enhance expression of pro-inflammatory chemokines such as CCL2, CCL5, and CXCL10, contributing to persistent inflammatory responses in skin tissues (*62*). By comparison, intact EGFR signaling can attenuate IFN-γ-induced pro-inflammatory gene programs, including chemokine expression, in epithelial cells (*63*). Thus, the exaggerated inflammatory responses observed during anti-EGFR therapy provide a clinically-relevant mirror image of the immunomodulatory effects observed following eAREG delivery. In addition, our findings may also provide a mechanistic framework linking Treg-derived growth factor signaling to the coordinated regulation of tissue regeneration and inflammation. AREG expression by Tregs in injured tissues may represent an endogenous strategy to simultaneously promote tissue healing while restraining excessive recruitment of inflammatory immune populations. In this context, AREG-mediated EGFR signaling in tissue-resident non-immune cells emerges as a mechanism that integrates morphogenic cues with negative feedback on chemokine-driven inflammation. By engineering AREG to enhance EGFR signaling while preserving its capacity for tissue retention, eAREG effectively amplifies this natural regulatory logic and translates it into a therapeutically effective strategy for promoting tissue healing under both physiological and pathological inflammatory conditions.

Importantly, the dual regenerative and immunomodulatory functions of eAREG were preserved in diabetic mice, a disease context characterized by high inflammation and profoundly impaired tissue healing. In fact, chronic non-healing skin wounds represent a major and growing clinical burden in patients with diabetes, frequently leading to infection, hospitalization, and limb amputation, with few effective regenerative therapies currently available (*64, 65*). Similarly, volumetric muscle loss injuries present a substantial unmet clinical need, arising not only from traumatic injury but also as a consequence of surgical procedures such as hip replacement, tumor resection, or severe orthopedic reconstruction, where endogenous regenerative capacity is insufficient to restore functional muscle tissue (*6, 66, 67*). In both settings, persistent inflammation is a key barrier to effective repair, and the ability of eAREG to simultaneously enhance tissue regeneration while suppressing pathological immune cell recruitment is therefore particularly relevant for these indications. By acting on tissue-resident cells to promote morphogenesis and restrain chemokine-driven inflammation, eAREG addresses two fundamental drivers of regenerative failure in chronic and complex injuries. Thus, these properties position eAREG as a promising therapeutic candidate for clinical contexts in which immune dysregulation actively undermines tissue repair.

In summary, this study identifies eAREG as a rationally engineered growth factor that integrates enhanced morphogenic signaling with active immunoregulatory control exerted via tissue-resident non-immune cells. By combining EGF-like signaling potency with ECM-mediated tissue retention and epigenetic suppression of inflammatory chemokine programs, eAREG promotes tissue healing through coordinated regenerative and immunomodulatory mechanisms. Beyond establishing eAREG as a promising therapeutic candidate, our findings also define a general design principle for regenerative biologics, in which growth factor potency, spatial persistence, and immune modulation are integrated to overcome barriers that limit tissue repair and regeneration in both healthy and disease contexts.

## Materials and Methods

### Study design

This study was designed to assess the regenerative and immunomodulatory functions of eAREG in *vivo* and *in vitro*. Murine models of excisional skin wounding and volumetric muscle loss were used to evaluate tissue repair, immune cell infiltration, and chemokine regulation following treatment with eAREG, recombinant AREG, EGF, or vehicle control. Animals were randomly allocated to treatment groups, and sample sizes were chosen based on prior studies using these models. For in vivo experiments, n denotes the number of independent wounds or muscle injuries analyzed per condition, with group sizes typically ranging from *n* = 6 to *n* = 10. Histological, microCT, and flow cytometric analyses were performed blinded to treatment condition where feasible. In vitro experiments were performed using independent biological replicates. Statistical analyses were conducted using predefined methods as described below.

### Mice and animal ethics

All animal studies were approved by the Monash Animal Research Platform Animal Ethics Committee, Monash University (approval number 17124). C57BL6/J mice were purchased from the Monash Animal Research Platform. *Lepr^db/db^* (BKS.Cg-*Dock7^m^+/+Lepr^db^*/J, strain 000642) were obtained from Jackson Laboratory. Mice were co-housed under a 12:12 light:dark light cycle at temperatures ranging between 20 and 24°C and humidities between 40 and 60%. Food and water were available *ad libitum* to the mice. Regular diet (BARASTOC) was provided. At any significant sign of distress or at the end of the experiments, mice were euthanized by cervical dislocation or CO_2_ asphyxiation.

### Skin wound healing model

C57BL6/J male mice aged 10–12 weeks or *Lepr^db/db^* aged 16–20 weeks were anaesthetized using isoflurane and given subcutaneous buprenorphine analgesia (0.1 mg/kg). Their backs were shaved, and two full-thickness excisional wounds were created using a 5 mm biopsy punch (Kai Medical). Each wound was splinted with an 8 mm nylon ring (M8 nylon washer, Zenith ITWProline) glued to the skin with Ultra-fast super glue (UHU). Rings were covered with small adhesive circular bandages (Elastoplast, Beiersdorf) and secured using 3M Blenderm surgical tape. Bandages and rings were inspected every alternate day and replaced if they started to loosen. This prevented wound healing via skin contraction, creating a model closer to human wound healing via re-epithelialization. Wounds were treated by intradermal injection of saline (PBS) or protein solutions in PBS. Wild-type mice received growth factors equimolar to a total of 1 µg AREG administered on days 1 and 4 post-wounding. Diabetic mice received eAREG at a total dose of 2 µg administered on days 1, 4, and 7 post-wounding. Solutions were delivered as four injections of 10 µl each around the edge of the wound.

### Volumetric muscle loss model

C57BL6/J male mice aged 10–12 weeks or *Lepr^db/db^* (BKS.Cg-*Dock7^m^+/+Lepr^db^*/J, Jackson Laboratory, strain 000642) aged 16–20 weeks were anaesthetized using isoflurane and given subcutaneous buprenorphine analgesia (0.1 mg/kg). The right hind limb was shaved, and a 1 cm unilateral incision exposed the underlying quadriceps and patella. The volume of muscle tissue undergoing volumetric muscle loss was determined as 7 mm-long from the patella’s outer edge (5 mm for diabetic mice) and affecting the rectus femoris, vastus medialis, and vastus lateralis muscles. Using scissors, the muscle mass as described above was excised. The vastus intermedius was not injured and the tendons were preserved. Then, the skin was closed with staples. Defects were covered with a fibrin matrix (60 µl total, 8 mg/ml fibrinogen (Enzyme Research Laboratories), 12 U/ml bovine thrombin (Sigma), 5 mM CaCl_2_, and 17 µg/ml aprotinin (Roche, Sigma)) containing growth factors equimolar to 4 µg of AREG or an equivalent volume of HEPES buffer.

### Histomorphometric analysis of skin wound closure

At day 7 or day 10 post-injury, mice were euthanized and the dorsal skin including the wounded area was excised. Wounds were fixed in 10% neutral buffered formalin for 24 hours at room temperature. Wounds were then harvested using an 8 mm biopsy punch around the original injury, embedded in paraffin, and sectioned into 4 µm thick sections until the center of the wound was passed. Sections were stained with haematoxylin and eosin and re-epithelialization was measured by histomorphometric analysis of tissue sections using Aperio ImageScope (Leica Biosystems). The center of wounds was determined by measuring the distance of the gap between the edges of the panniculus carnosus muscle that is severed upon wound creation. Wound closure was calculated, as the ratio of epidermis closure to the length of the panniculus carnosus gap.

### Histomorphometric analysis of muscle regeneration

At day 21 after the injury, mice were euthanized, limbs were collected and fixed in 10% formalin for 24 hours at room temperature and then decalcified with 5% nitric acid overnight. The quadriceps muscles, together with the femur and the patella, were removed and embedded in paraffin. Four samples per group were sliced into 4 µm thick longitudinal sections. Sections were stained with Masson’s Trichrome, and images were analyzed using Aperio ImageScope software (Leica Biosystems). The injured area was defined as 7 mm-long from the patella’s outer edge, and above the patella’s superior edge. Fibrosis percentage was calculated as the area of fibrosis tissue divided by the area of muscle on sections representative of the longitudinal middle of the quadriceps.

### MicroCT

Fixed limbs were scanned with an Inveon micro-CT (Siemens) operated at an energy of 80 kV, 500 µA. Scans were reconstructed with a Feldkamp algorithm at a nominal isotropic resolution of 30 µm. After reconstruction, images were rotated to align the longitudinal axis of the femur with the axes of the images. A cubic mask (7 x 7 x 7 mm for wild-type mice and 5 x 7 x 7 mm for diabetic mice) was applied over the injured area. Muscle volume was calculated by applying a visual threshold of - 200/+400 HU using the IRW software. Volume of tissue restoration was calculated as the ratio of muscle volume between the injured leg and the uninjured leg.

### Recombinant protein production and purification

Sequences were cloned into the expression vector pET-22b(+) (GE Healthcare) and expressed in *Escherichia coli* BL21 (DE3). A 6×His tag was added at the N-terminus of the eAREG sequence. Bacteria were cultured overnight in 10 ml of lysogeny broth (LB) containing 100 µg/ml ampicillin. The overnight culture was diluted 1:100 into fresh LB medium supplemented with 100 µg/ml ampicillin and 100 µg/ml carbenicillin and grown at 37°C for 3 hours. Protein expression was induced with 1 mM isopropyl β-D-1-thiogalactopyranoside (IPTG), and cultures were incubated overnight at 30°C. Cells were harvested by centrifugation at 4,000 x g for 10 minutes, and the pellet was resuspended in cold lysis buffer containing 2 mM 2-mercaptoethanol (Sigma-Aldrich), 0.2 mg/ml lysozyme (Roche), 1% Triton X-100, 1% Zwittergent (Sigma-Aldrich), 10 mM MgCl₂, and a protease inhibitor cocktail tablet (Roche, 11836170001) in Tris-EDTA buffer supplemented with NaCl (pH 8.0). Cells were lysed by sonication for 30 seconds at maximum amplitude for four cycles. Benzonase (500 U; Millipore) was added, and lysates were incubated on a rotating platform for 1 hour at 4°C. Lysates were centrifuged at 13,000 x g for 15 minutes, and the supernatant was discarded. The pellet was resuspended in cold extraction buffer containing 6 M guanidine hydrochloride (GuHCl), a protease inhibitor cocktail tablet, and CAPS buffer, followed by incubation on a rotating platform for 15 minutes at 4°C. The extract was centrifuged again at 13,000 x g for 15 minutes, and the supernatant was filtered through a 0.22 µm filter. Proteins were purified using a HisTrap HP 5 ml affinity column (GE Healthcare) under denaturing conditions and refolded on-column using a linear gradient from 100% to 0% GuHCl. Chaperone proteins were removed by washing with ATP buffer (50 mM Tris-HCl, 150 mM NaCl, 10 mM MgSO_4_, 2 mM ATP, pH 7.4). Lipopolysaccharides were removed using Triton X-114 phase separation (PBS containing 0.1% Triton X-114). Proteins were further purified by size-exclusion chromatography (SEC; GE Healthcare). The final protein preparations were dialyzed against HEPES buffer (20 mM HEPES, 150 mM NaCl), sterile-filtered through a 0.22 µm filter, and stored at −80°C. Protein purity (>99%) was confirmed by SDS-PAGE. Endotoxin levels were measured using the Pierce Chromogenic Endotoxin Quant Kit (Thermo Fisher Scientific, A39552S), and only samples containing <0.2 EU/µg endotoxin were used.

### Commercial proteins

The following recombinant proteins were purchased: EGFR (Abcam, ab167752); AREG (Peprotech, 315-36); EGF (R&D Systems, 2028EG); HB-EGF (ab215635) tenascin-C (Merck, CC065); collagen type I (Merck, CC050); fibronectin (Sigma-Aldrich, F2006); osteopontin (PeproTech, 120-35); vitronectin (PeproTech, 140-09); fibrinogen (Enzyme Research Laboratories, FIB3).

### Growth factor binding to ECM proteins

ELISA plates (Greiner Bio-one medium binding, Thermo Fisher Scientific) were coated with 100 nM of ECM protein (vitronectin, tenascin-C, fibrinogen, fibronectin, osteopontin, collagen II) for 1 hour at 37°C and blocked with PBS containing 1% BSA (Sigma-Aldrich, A7030) for 1 hour at room temperature. Then, plates were washed three times with PBS-T (0.2% Tween-20, pH 7.2) and incubated 1 hour at room temperature with growth factors (200 nM in PBS with 0.1% BSA). Plates were washed three times with PBS-T and growth factors were detected with biotinylated anti-EGF detection antibody (R&D Systems, DY2028) in PBS-T with 0.1% BSA (2 hours at room temperature). Plates were washed three times with PBS-T and biotin was detected with HRP-conjugated streptavidin (2 µg/ml in PBS-T, 20 minutes at room temperature). Wells were washed three times with PBS-T and detection was done with tetramethylbenzidine substrate and measurement of the absorbance at 450 nm.

### Growth factor binding to heparan sulfate

Heparin-binding plates (Corning, 354676) were coated with 25 µg/ml of heparan sulfate (Sigma-Aldrich, H7640) overnight at room temperature and blocked with a PBS solution containing 0.2% gelatin and 0.5% BSA for 1 hour at room temperature. Wells were washed 3 times with washing buffer (100 mM NaCl, 50 mM NaAc, 0.2% Tween-20, pH 7.2) and growth factors (200 nM in PBS with 0.5% BSA) were added. After 1 hour incubation at room temperature, wells were washed three times with the washing buffer and bound growth factors were detected with biotinylated anti-EGF detection antibody (R&D Systems, DY2028) or anti-histidine tag (Abcam, ab27025) in PBS-T with 0.1% BSA (2 hours at room temperature). Plates were washed three times with PBS-T and biotin was detected with HRP-conjugated streptavidin (2 µg/ml in PBS-T, 20 minutes at room temperature). Wells were washed three times with PBS-T and detection was done with tetramethylbenzidine substrate and measurement of the absorbance at 450 nm.

### Growth factor binding to EGFR

ELISA plates (Greiner Bio-one medium binding, Thermo Fisher Scientific) were coated with 50 nM of EGFR overnight in PBS at 4°C and blocked with PBS containing 2% BSA (Sigma-Aldrich, A7030) for 1 hour at room temperature. Plates were washed three times in PBS-T (0.2% Tween-20, pH 7.2) and AREG, EGF, or eAREG were added in serial two-fold dilutions ranging from 100 nM to 0 nM in PBS-T with 1% BSA (1 hour at room temperature). Plates were washed three times with PBS-T and EGF was detected using biotinylated anti-EGF detection antibody (R&D Systems, DY2028), while AREG and eAREG were detected using biotinylated anti-AREG antibodies (R&D Systems, BAF989) in PBS-T containing 1% BSA (2 hours at room temperature). Plates were washed three times with PBS-T and biotin was detected with HRP-conjugated streptavidin (2 µg/ml in PBS-T, 20 minutes at room temperature). Wells were washed three times with PBS-T and detection was done with tetramethylbenzidine substrate and measurement of the absorbance at 450 nm. The dissociation constant (*K_d_*) was obtained by fitting the binding curves with a specific binding with Hill Slope equation (Prism, Version 10, GraphPad, USA).

### Cells

Immortalized human keratinocyte cells (HaCaT, gifted by Professor Richard Boyd, Monash University) were maintained in Roswell Park Memorial Institute 1640 medium (RPMI, Thermo Fisher Scientific) supplemented with 10% fetal bovine serum (FBS, Thermo Fisher Scientific). Human myoblasts derived from primary skeletal muscle satellite cells (Lifeline Cell Technology, FC-0091) were maintained in StemLife Sk Complete Medium (Lifeline Cell Technology, LL-0069). Primary mouse fibroblasts from C57BL/6J mouse tails were isolate and maintained as described previously (*20*). Primary mouse MSCs from C57BL6/J mice were isolated and maintained as described previously (*27*). All cell lines were maintained in culture with 100 µg/ml streptomycin and 100 units/ml penicillin at 37°C with 5% CO_2_.

### EGFR internalization and degradation

HaCaT cells (50,000 per well) were seeded in 24-well plates and serum-starved in low-serum medium (RPMI containing 0.1% FBS) for 24 hours. Cells were then treated with growth factors at equimolar concentrations corresponding to 100 ng/ml EGF for 15, 30, 60, 120, 180, or 360 minutes, or with an equivalent volume of PBS to determine basal levels of receptor turnover. Following stimulation, cells were washed with acidic glycine buffer (50 mM glycine, 150 mM NaCl, pH 2.7) for 1 minute at room temperature to remove growth factors bound to EGFR. Cells were subsequently detached using TrypLE (Thermo Fisher Scientific) supplemented with 20 mM EDTA and stained with a fixable viability dye (LIVE/DEAD Aqua, Thermo Fisher Scientific, L34957) for 30 minutes on ice. Samples were then split into two fractions and processed as follows. To assess EGFR internalization, surface EGFR was stained using an anti-human EGFR antibody (BioLegend, clone AY13, 2 µg/ml) in FACS buffer (PBS containing 5% FBS and 2 mM EDTA) for 30 minutes on ice. To assess EGFR degradation, cells were fixed and permeabilized (eBioscience, 00-5523-00) for 30 minutes on ice, followed by intracellular staining with the same anti-human EGFR antibody for 30 minutes on ice. Samples were acquired on a flow cytometer (Cyan, Beckman Coulter) and analyzed using FlowJo software (version 11).

### Phosphorylation array

HaCaT cells (30,000 per well) were plated in 24-well plates and serum-starved in low-serum medium (RPMI containing 0.1% FBS) for 24 hours. Cells were then treated with growth factors at equimolar concentrations corresponding to 100 ng/ml EGF for 10, 30, 180, or 360 minutes, or with an equivalent volume of PBS to assess basal receptor phosphorylation. Following stimulation, cells were washed with ice-cold PBS, and samples were prepared according to the manufacturer’s instructions for the Human Phospho-EGFR Phosphorylation Antibody Array kit (Abcam, ab134005). Membranes were imaged using a high-sensitivity CCD camera system (ChemiDoc Touch Imaging System, Bio-Rad). Signal intensity was quantified by densitometry using Image Lab software (Bio-Rad). Relative phosphorylation levels were calculated according to the manufacturer’s instructions and normalized to both the membrane positive controls and untreated control samples (low-serum medium only).

### EGFR and ERBB2 total phosphorylation

HaCaT cells (30,000 per well) were seeded in 24-well plates and serum-starved in low-serum medium (RPMI containing 0.1% FBS) for 24 hours. Cells were then treated with growth factors at equimolar concentrations corresponding to 100 ng/ml EGF for 10, 30, 180, or 360 minutes, or with an equivalent volume of PBS to assess basal receptor phosphorylation. Following stimulation, cells were washed with ice-cold PBS, and samples were processed according to the manufacturer’s instructions using either the Human Phospho-EGFR DuoSet IC ELISA (Abcam, DYC1095B) or the Human Phospho-ErbB2/HER2 DuoSet IC ELISA (Abcam, DYC1768). Total receptor phosphorylation was quantified as specified by the manufacturer.

### Proliferation assay

Cells were starved for 24 hours (respective media with 2% FBS) and then seeded on a 96-well cell culture plate (2,000 cells/well for HaCaT; 1,000 cells/well for fibroblasts, myoblast, and MSCs) in combination with growth factor (equimolar to 20 ng/ml EGF) in low-serum media (2% FBS) for 72 hours. Percentage of new cells was calculated over basal proliferation (low-serum media only) using CyQuant (Thermo Fisher Scientific, C7026) and the equation [(cell number in basal proliferation group/cell number in stimulation group) – 1] × 100.

### Migration assay

Assays were conducted using 6.5-mm-diameter culture plate inserts (Corning, 3421) with 5 µm pore sizes. Cells (1 x 10^5^) in migration media (DMEM/F12 with 0.25% BSA) were added to the inserts. The lower chambers contained migration buffer alone or growth factors (equimolar to 20 ng/ml EGF). Cells were allowed to migrate through the insert membrane for 6 hours at 37°C with 5% CO_2_. The inserts were then fixed with 4% paraformaldehyde, and cells on the upper side were removed. DAPI (1 µg/ml) was used to stain cells on the bottom side, and they were counted using a fluorescent microscope. The data are presented as the fold change, calculated by dividing the number of cells that migrated in response to treatments by the number of cells that migrated spontaneously (migration media only).

### Flow cytometry

Skin wounds were harvested using an 8 mm biopsy punch, and muscle defects were dissected to isolate the quadriceps. Samples were minced with scissors and subjected to two serial digestions with collagenase XI (1 mg/ml) at 37°C (30 minutes for skin, 20 minutes for muscle). After the first digestion, the supernatant was collected and mixed with neutralisation buffer (DMEM/F12 with 10% FBS and 5 mM EDTA). The first collection was kept on ice, and fresh collagenase XI was added to the undigested tissue for the second digestion. Digestion mixtures were passed through a 70 µm cell strainer and stained with LIVE/DEAD Fixable Aqua dye (Thermo Fisher Scientific, 1:400 dilution in PBS) for 20 minutes on ice. Cells were incubated with TruStain FcX anti-CD16/32 (10 µg/ml; clone 93, BioLegend) diluted in staining buffer (5% FBS and 2 mM EDTA in PBS) for 20 minutes and subsequently incubated with primary antibodies in staining buffer for a further 30 minutes on ice. The following anti-mouse antibodies from BioLegend were used: FITC anti-CD11b (clone M1/70, 6.6 μg/ml) or BV711 anti-CD11b (clone M1/70, 2 µg/ml); PE anti-F4/80 (clone BM8, 4 µg/ml); BV421 anti-Ly6G (clone 1A8, 2 µg/ml); BV711 anti-Ly6C (clone HK1.4, 1 μg/ml) or FITC anti-Ly6C (clone HK1.4, 5 µg/ml); PE-Cyanine7 anti-CD206 (clone C068C2, 2.6 µg/ml); PE-Cyanine7 anti-CD3 (clone 17A2, 4 µg/ml); APC anti-CD4 (clone GK1.5, 2 μg/ml); BV421 anti-CD8 (clone 53-6.7, 2 µg/ml); APC/Fire 750 anti-TCR γ/δ (clone GL3, 2 µg/ml); FITC anti-CD19 (clone 1D3, 1 µg/ml); and BV421 anti-NK-1.1 (clone PK136, 1 µg/ml). Cells were washed once with a large volume of staining buffer before analysis with BD LSR.

### Growth factor retention

C57BL/6 mice (10-week-old) were used. For retention in skin wounds, full-thickness punch-biopsy wounds (5 mm in diameter) were created. Then, 1 µg of AREG or an equimolar amount of eAREG and EGF was injected intradermally in four sites around the wound area (10 µl per injection). Full-thickness skin tissue around the wound was harvested with an 8 mm biopsy punch at day 1, 3, or 5 post-injections. For retention experiments in quadricep muscle defects, 4 µg of AREG or an equimolar amount of eAREG and EGF was delivered using a fibrin matrix. Then, the entire quadriceps containing the fibrin matrix was harvested at day 1, 3, and 5, and 7 post-delivery. All harvested tissues were transferred into 500 µl of T-PER Tissue Protein Extraction Reagent (Thermo Fisher Scientific, 78510) containing a protease inhibitor cocktail (1 tablet for 50 ml, Roche, 11836170001) and minced. Samples were incubated for 1 hour at room temperature under agitation, centrifuged at 5,000 x g for 5 minutes and supernatants were stored at -80°C. Concentration of proteins were determined by ELISA using an anti-OLLAS tag capture antibody (1 µg/ml, Novus Biologicals, NBP1-06713) and a detection antibody from AREG or EGF DuoSet ELISA kit (DY989 or DY2028, R&D Systems). Day 0 samples were collected right after delivery and used to make a standard curve and to determine the 100% value.

### Expression of Egfr and Erbb2 in injured tissues

Mouse skin wounds and injured muscle tissue were harvested on day 3 post-injury and digested as described for flow cytometry. Dissociated cells were stained with TruStain FcX anti-mouse CD16/32 antibody (10 µg/ml; clone 93, BioLegend), followed by cell surface staining with antibodies from BioLegend (FITC anti-mouse CD45, clone 30-F11, 6 µg/ml; and APC-Fire anti-mouse CD11b, clone M1//70, 6 µg/ml), diluted in FACS buffer. Cell viability was detected with 4′, 6-diamidino-2-phenylindole dihydrochloride (DAPI,1 µg/ml, Sigma). Cells were sorted into non-hematopoietic (CD45^-^), myeloid (CD45^+^CD11b^+^), and non-myeloid (CD45^+^CD11b^-^) cell populations using a BD Influx Cell Sorter (BD Biosciences). Sorted cells were collected, and RNA was extracted using the Quick-RNA MicroPrep Kit (Zymo Research) to detect *Egfr* and *Erbb2* expression. Reverse transcription and quantitative PCR were performed using SuperScript IV Reverse Transcriptase (ThermoFisher), and ThermoFisher TaqMan Assay primers (egfr, Mm01187858_m1; erbb2, Mm00658541_m1; Actb, Mm02619580_g1; Gapdh, Mm99999915_g1), on a LightCycler96 instrument with software LightCycler 96 (Roche Diagnostics). Relative gene expression levels were calculated using the 2(^-ΔCt^) method, normalized to the reference gene (*Actb* and *Gapdh*).

### Chemokine array

Keratinocytes, myoblasts, fibroblasts and MSCs were cultured as described above in 6-well plates containing 2 ml of media per well until reaching 70% confluency. Media was then removed, cells were rinsed with PBS, and 2 ml of low-serum media (respective basal media + 2% FBS and 1% penicillin/streptomycin) was added per well. Cells were incubated for a further 24 hours at 37°C with 5% CO_2_. Cells were then treated with 4 ng/ml recombinant human or mouse TNF-α (PeproTech), 1 ng/mL recombinant human or mouse IL-1β (PeproTech), 0.1 ng/ml recombinant human or mouse interferon-γ (PeproTech), and 2 µg/ml polymyxin B (Sigma-Aldrich), with or without 20 ng/ml eAREG, all prepared in PBS (Sigma-Aldrich). Negative control cells were treated with an equivalent volume of PBS and 2 µg/ml polymyxin B. eAREG only control cells were treated with 20ng/ml eAREG and 2 µg/ml polymyxin B. After a further 24 hours, media were collected, centrifuged to remove cells and debris, and supernatants processed for analysis using Proteome Profiler Human or Mouse Chemokine Array Kits (R&D Systems) according to the manufacturer’s instructions. Membranes were imaged using the G:Box (Syngene) gel dock. Densitometry analysis to compare levels of detected chemokines between conditions was performed using ImageJ.

### Chemokine ELISAs

Cells were treated as described for the chemokine array assay. Chemokine concentrations in culture supernatants were quantified using DuoSet ELISA kits (R&D Systems): Human CCL2/MCP-1 (DY279), Mouse CCL2/JE/MCP-1 (DY479), Human CCL5/RANTES (DY278), Mouse CCL5/RANTES (DY478), Human CXCL9/MIG (DY392), Mouse CXCL9/MIG (DY492), Human CXCL10/IP-10 (DY266), and Mouse CXCL10/IP-10/CRG-2 (DY466).

### ATAC-seq

Keratinocytes cells were cultured at 37°C with 5% CO_2_ in 6-well plates containing 2 ml RPMI-1640 (Thermo Fisher Scientific) supplemented with 10% fetal bovine serum (FBS) (Thermo Fisher Scientific) and 1% penicillin/streptomycin (Thermo Fisher Scientific) per well until reaching 70% confluency. Media was then removed, cells were rinsed with PBS, and 2 ml of low-serum media (RPMI-1640 supplemented with 2% FBS and 1% penicillin/streptomycin) was added per well. Cells were incubated for a further 24 hours at 37°C with 5% CO_2_. Cells were then treated with 4 ng/ml recombinant human TNF-α (PeproTech), 1 ng/mL recombinant human IL-1β (PeproTech), 0.1 ng/ml recombinant human interferon-γ (PeproTech), and 2 µg/ml polymyxin B (Sigma-Aldrich), with or without 20 ng/ml eAREG, all prepared in PBS (Sigma-Aldrich). Negative control cells were treated with an equivalent volume of PBS and 2 µg/ml polymyxin B. These conditions were selected to assess inflammation-induced chromatin accessibility, as epigenetic repression at promoter and enhancer regions is a known mechanism by which EGFR signaling regulates chemokine gene expression (*53*). After 24 hours, media was removed and cells were rinsed with PBS, then collected by trypsinization using TrypLE (Thermo Fisher Scientific). Trypsin was neutralized with RPMI-1640 containing 10% FBS, and cells were centrifuged at 300 x g for 5 minutes. The supernatant was removed, cells were resuspended in PBS to remove residual media, centrifuged again at 300 x g for 5 minutes, and resuspended in fresh PBS for counting. A total of 50,000 cells per condition were transferred to 1.5 mL tubes, and ATAC-seq libraries were prepared using the Zymo-Seq ATAC Library Kit (Zymo Research) according to the manufacturer’s instructions. Libraries were purified using AMPure XP double-sided purification (Beckman Coulter) and assessed by Qubit fluorometry (Thermo Fisher Scientific), capillary electrophoresis, and quantitative PCR. For sequencing, library pools were loaded at 800 pM with approximately 1% PhiX for onboard denaturation and clustering. Sequencing was performed using a P3 100-cycle kit on a NextSeq 2000 instrument (Illumina). Base calling and demultiplexing were performed using DRAGEN BCL Convert version 4.2.7 (Illumina). Raw sequencing data were processed using the nf-core/atacseq pipeline (version 2.1.2) (*68*) with the GRCh38 Ensembl genome assembly (release 95) for quality control, alignment, and peak calling. Peak visualization and inspection were performed using Integrative Genomics Viewer version 2.19.4 with the GRCh38 (hg38) genome assembly. Peak annotations were generated using HOMER as implemented in the nf-core/atacseq pipeline or assessed manually using annotations from the GeneHancer database (*69*).

### Detection of chemokines *in vivo*

Homogenized skin wound and muscle tissues were incubated for 30 minutes on ice in T-PER Tissue Protein Extraction Reagent (10 ml/g tissue; Thermo Fisher Scientific) supplemented with protease inhibitor (1 tablet per 7 ml; Roche). Samples were then centrifuged at 10,000 x g for 5 minutes, and supernatants were collected and stored at −80°C. Total protein concentration was determined using a Bradford assay (Millipore). Chemokine levels were quantified using DuoSet ELISA kits (R&D Systems): Mouse CCL2/JE/MCP-1 (DY479), Mouse CCL5/RANTES (DY478), Mouse CXCL9/MIG (DY492), and Mouse CXCL10/IP-10/CRG-2 (DY466). Chemokine concentrations were normalized to total protein content and expressed as chemokine amount per mg of total protein.

### Statistical analysis

Statistical analyses were performed using GraphPad Prism 10 statistical software (GraphPad, USA). Significant differences were calculated with Student’s *t*-test, one-sample *t*-test, and by analysis of variance (ANOVA) when performing multiple comparisons between groups. *P* < 0.05 was considered statistically significant.

## Acknowledgments

We thank the Monash Histology Platform for assistance with histology, Adele Barugahare at the Monash Genomics Bioinformatics Platform for assistance with ATAC sequencing analysis, and Dr Michael De Veer at the Monash Biomedical Imaging Platform for assistance with microCT. Biorender.com was used to create some illustrations.

## Author contributions

M.M.M. conceptualized the initial study. Z.J. and M.M.M. designed the protein engineering strategy. J.M.D.L., C.P., Y-Z.L., and M.M.M. designed the experiments. J.M.D.L., C.P., Y-Z.L., B.N., Y.L., S.L., J.A.L., T.W., Z.J., and M.M.M performed the experiments and analysed the data. J.M.D.L., C.P., and M.M.M. wrote the manuscript.

## Data and materials availability

All data and code needed to evaluate and reproduce the results in the paper are present in the paper and/or the Supplementary Materials. ATAC-seq data have been deposited in the Gene Expression Omnibus (GEO) under accession number GSE328352 (https://www.ncbi.nlm.nih.gov/geo/query/acc.cgi?acc=GSE328352). Details of eAREG production are provided in the Methods section, Protein production and purification.

## Supplementary Materials

**Supplementary Figure 1.**
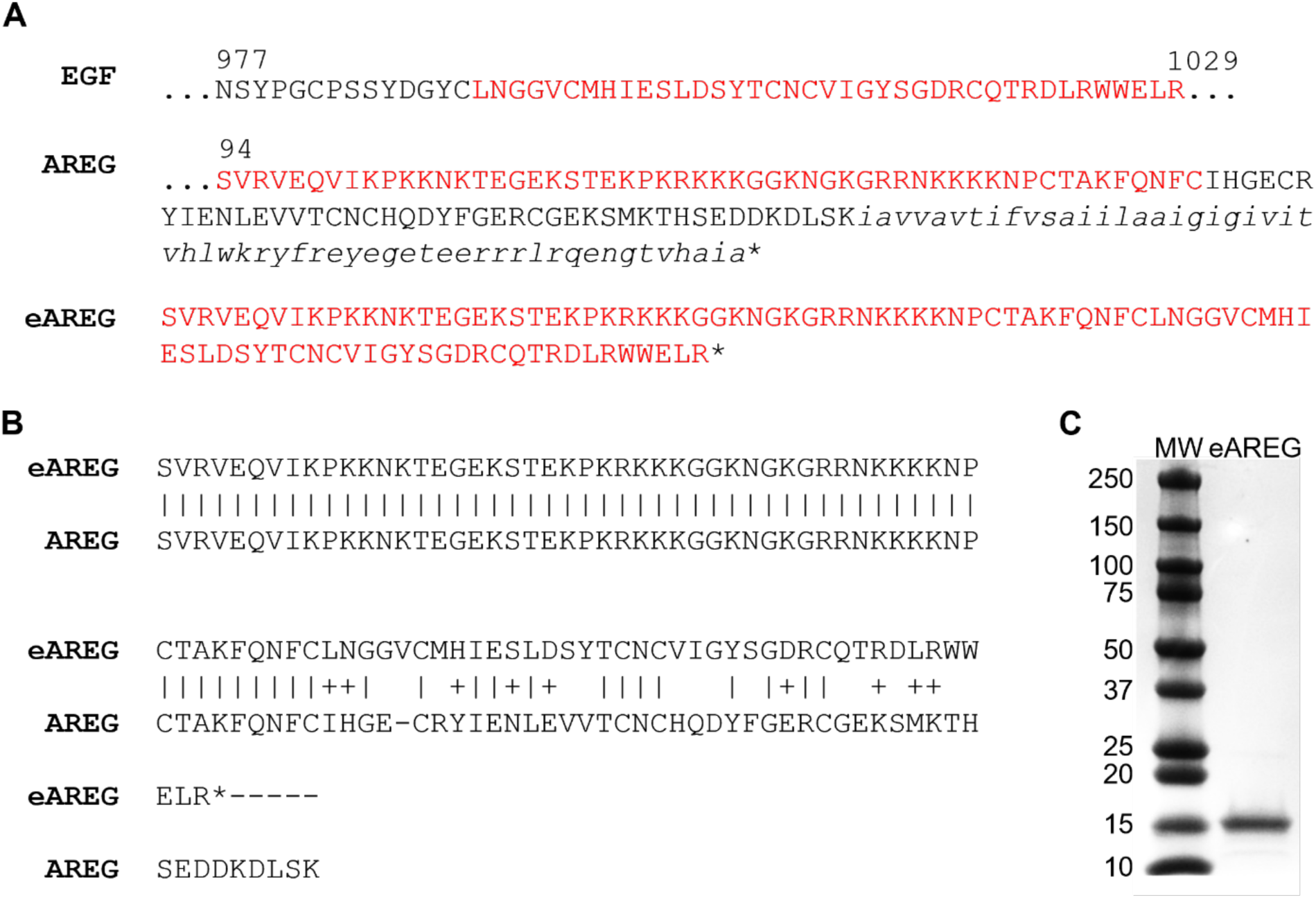
Design of eAREG which contains EGF- and AREG-derived domains. (**A**) Mature engineered AREG (eAREG) retains the N-terminal heparin-binding domain of AREG (red text in AREG sequence) and its EGF-like domain from the second cysteine residue (black, non-italicized text in AREG sequence) has been substituted for the EGFR-binding domain of EGF, also from its second cysteine residue (red text in EGF sequence). * indicates the protein C-terminus. The italicized text represents the AREG transmembrane domain that is proteolytically processed and not present in the mature, secreted protein. (**B**) Sequence alignment using protein BLAST demonstrates that eAREG is 76% identical to AREG. (**C**) Post-purification SDS-PAGE results for recombinant eAREG expressed in *Escherichia coli* demonstrate it is readily produced at high purity. MW indicates molecular weight markers, with weights indicated in kilodaltons.

**Supplementary Figure 2.**
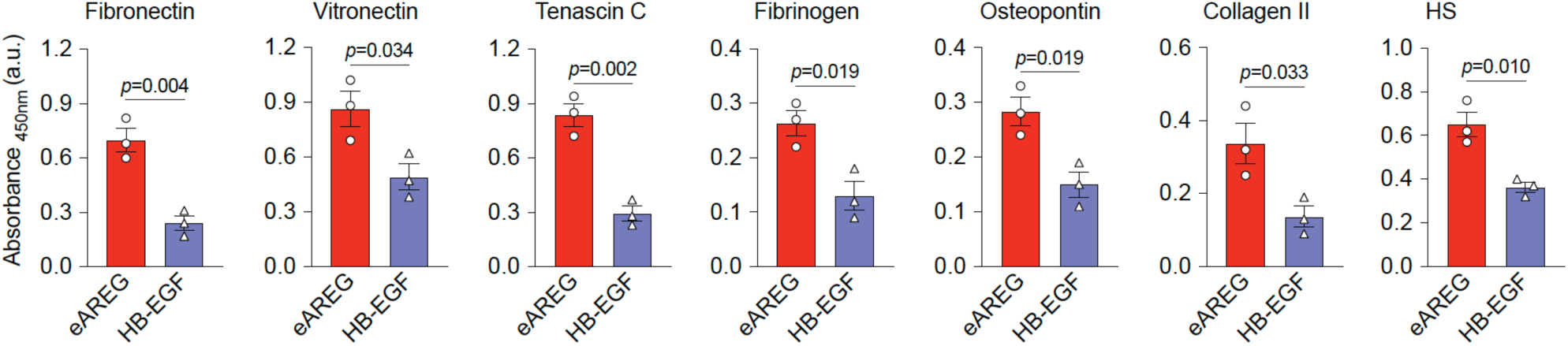
eAREG shows stronger binding to ECM components compared to HB-EGF. ELISA plates were coated with ECM proteins or heparan sulfate (HS) and incubated with eAREG or HB-EGF (200 nM equivalent to AREG). Bound eAREG and HB-EGF were detected with an antibody recognizing His-tag. *n* = 3 independent experiments. Statistical significance was determined using two-tailed unpaired Student’s *t* test. *P* values are indicated.

**Supplementary Figure 3.**
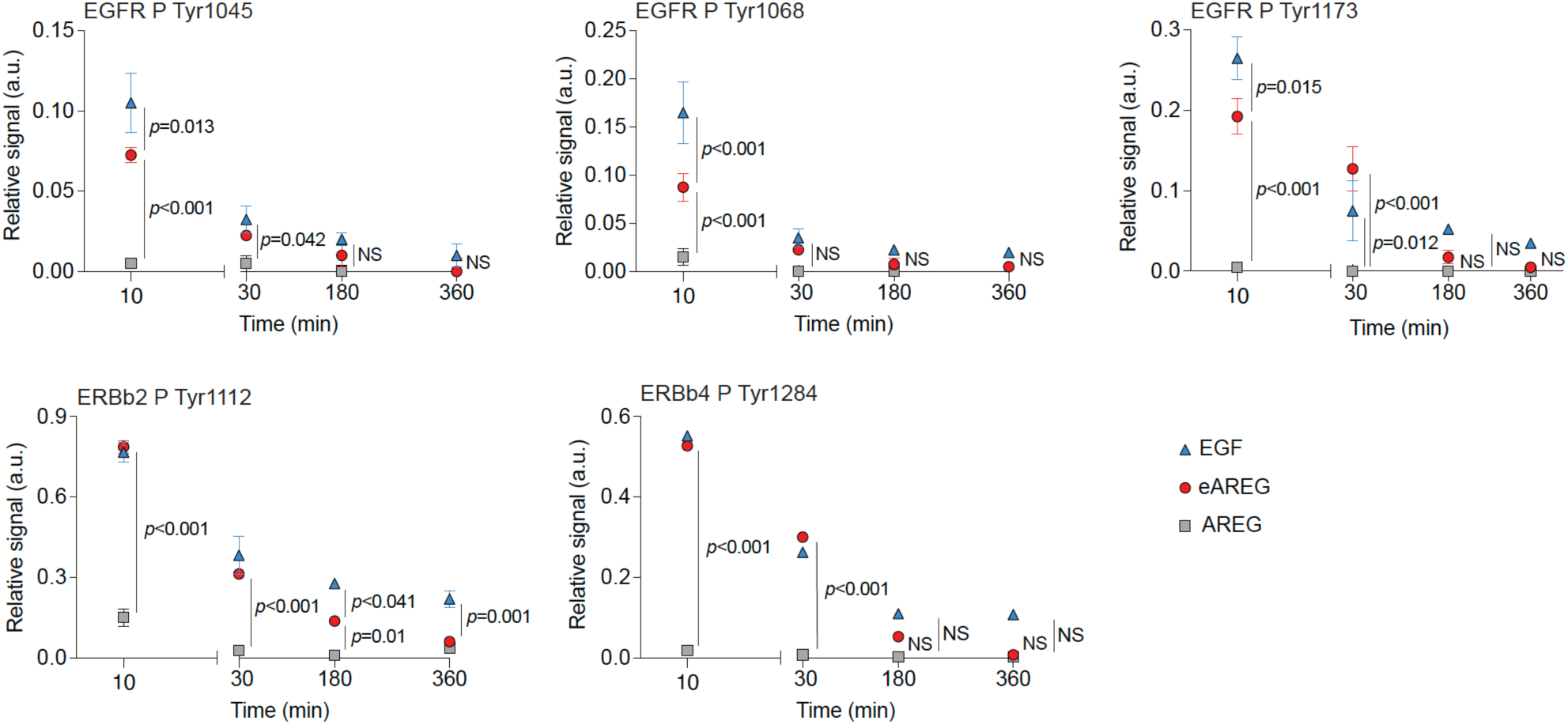
eAREG induces EGF-like phosphorylation of EGFR family receptors. Keratinocytes were treated with equimolar concentrations of EGF, AREG or eAREG and phosphorylation of specific tyrosine residues (P Tyr; residue number indicated) of EGFR family receptors (EGFR, ERBB2 and ERBB4) was quantified over time using antibody arrays. *n* = 3 independent experiments. Data are presented as mean ± SEM. Statistical significance was determined using two-way ANOVA with Bonferroni post hoc test for multiple comparisons. *P* values are indicated. NS, not significant.

**Supplementary Figure 4.**
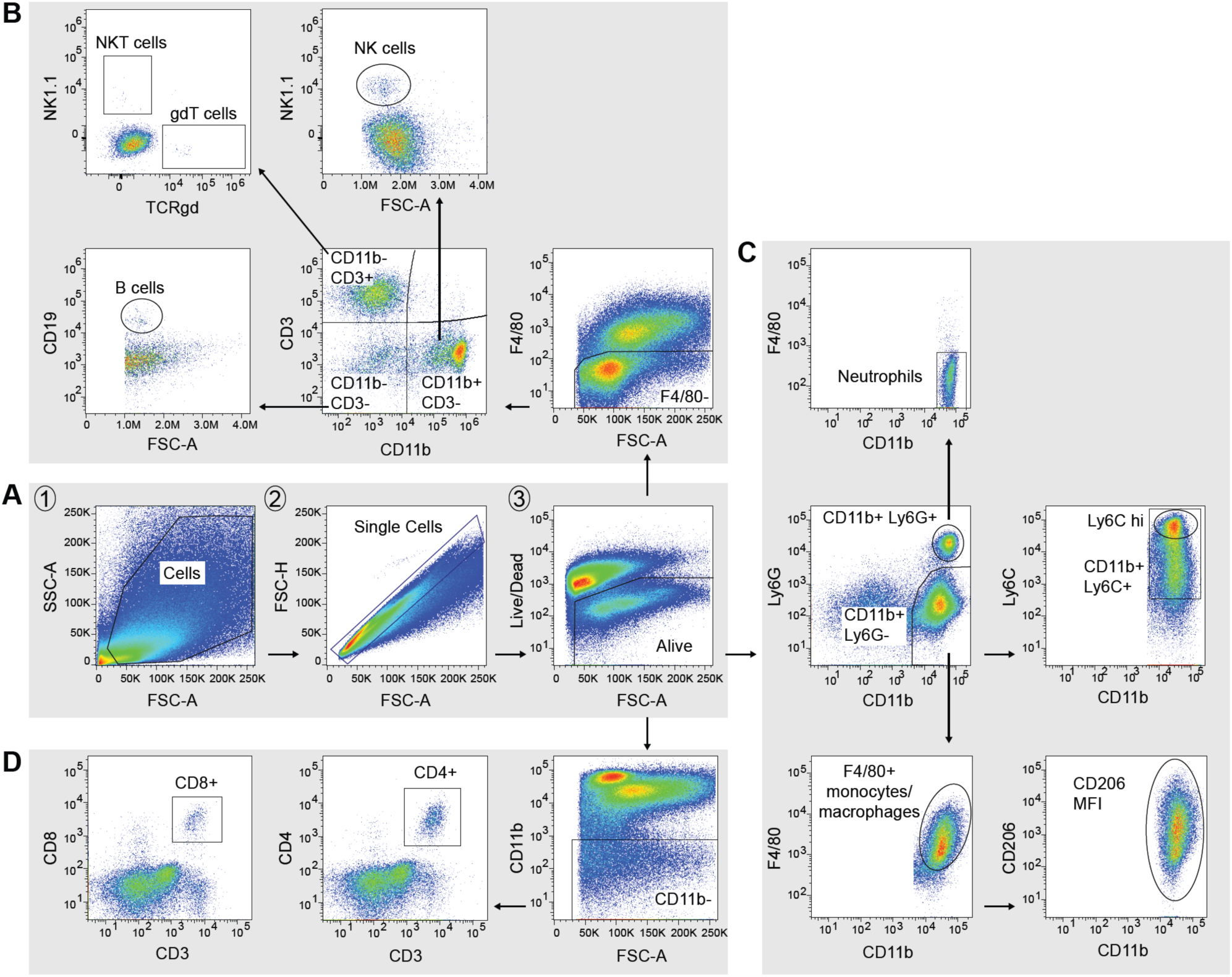
Representative flow cytometry gating strategies for analyzing immune cell subsets. (**A**) Sequential gating strategy to exclude debris, doublets, and dead cells, applied as the initial gating steps across all analyses. (**B**) Subsequent gating of lymphoid and myeloid cell subsets, where F4/80^-^ cells were separated into four fractions based on CD11b and CD3 expression, then further defined into CD19⁺ B cells, NK1.1⁺ NK and NKT cells, and TCRγδ⁺ γδ T cells. (**C**) Gating strategy for myeloid populations, including CD11b⁺ Ly6G⁺ F4/80⁻ neutrophils, and CD11b⁺ Ly6G⁻ Mo/MΦ, which was further subdivided into F4/80^+^ and Ly6C⁺ Mo/MΦ subsets. The CD11b⁺ F4/80⁺ Mo/MΦ were analyzed for CD206 expression by measuring the median fluorescence intensity (MFI). (**D**) T cells were gated by excluding CD11b⁺ myeloid cells, followed by identification of CD3⁺ CD8⁺ and CD3⁺ CD4⁺ T cell subsets.

**Supplementary Figure 5.**
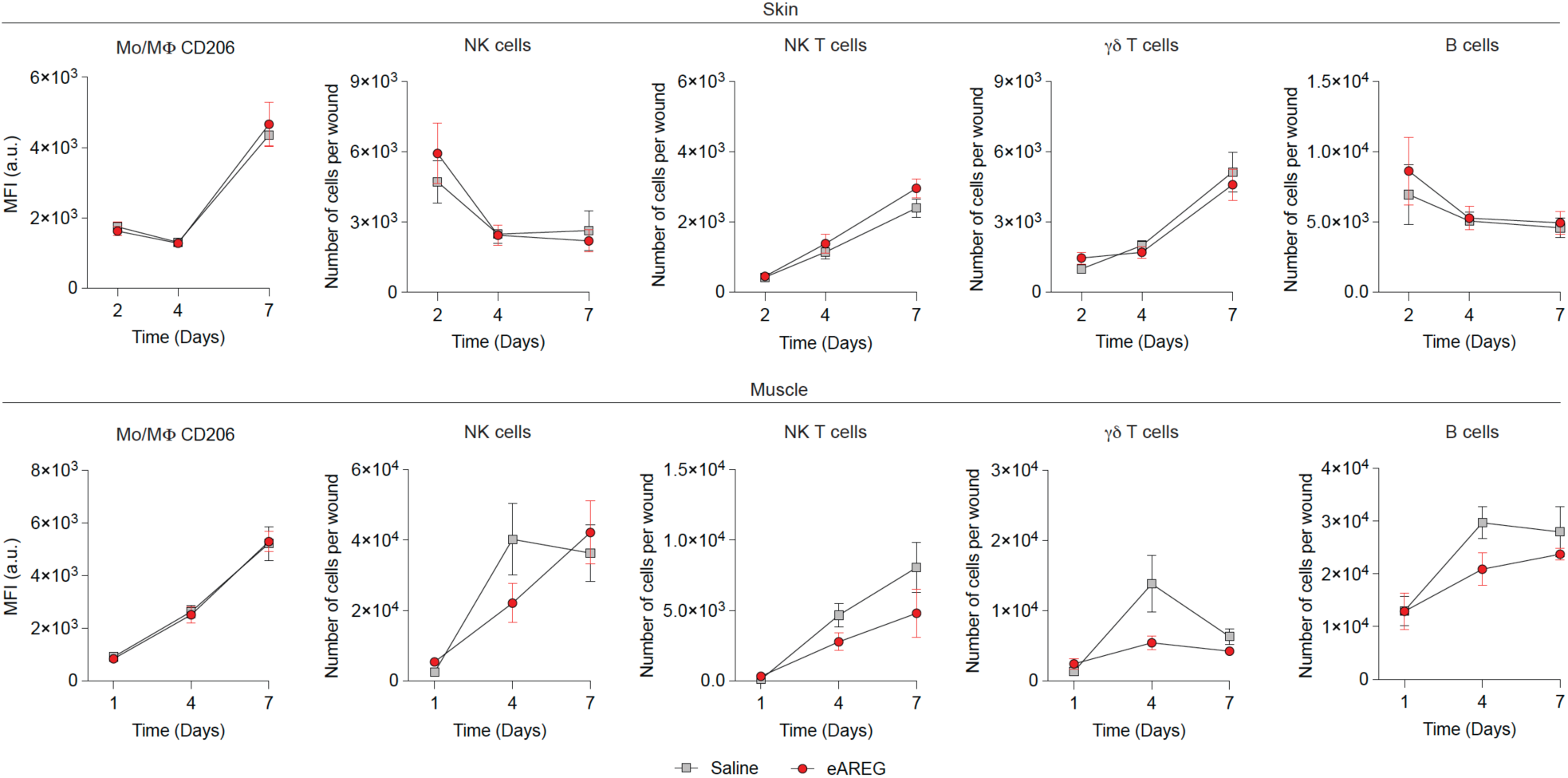
Delivery of eAREG does not influence NK cell, NKT cell, γd T cell, and B cell accumulation in injured tissues. Analysis of natural killer cells (NK cells), natural killer T cells (NK T cells), γδ T cells, and B cells populations by flow cytometry during tissue healing in wild-type mice. MFI of CD206 in Mo/MΦ was used to assess M2-like polarization. Skin wounds were treated with saline or eAREG at day 1 and 4 post-wounding and analyzed at day 2, 4, and 7 post-injury. Muscle injuries were treated with saline or eAREG and analyzed at day 1, 4, and 7 post-injury. *n* = 10 skin wounds and *n* = 8 muscle injuries per time point. Data are plotted as kinetic line plots showing mean ± SEM. Statistical significance was determined using two-way ANOVA with Bonferroni post hoc tests. No statistically significant differences were observed between conditions for any of the analyzed populations.

**Supplementary Figure 6.**
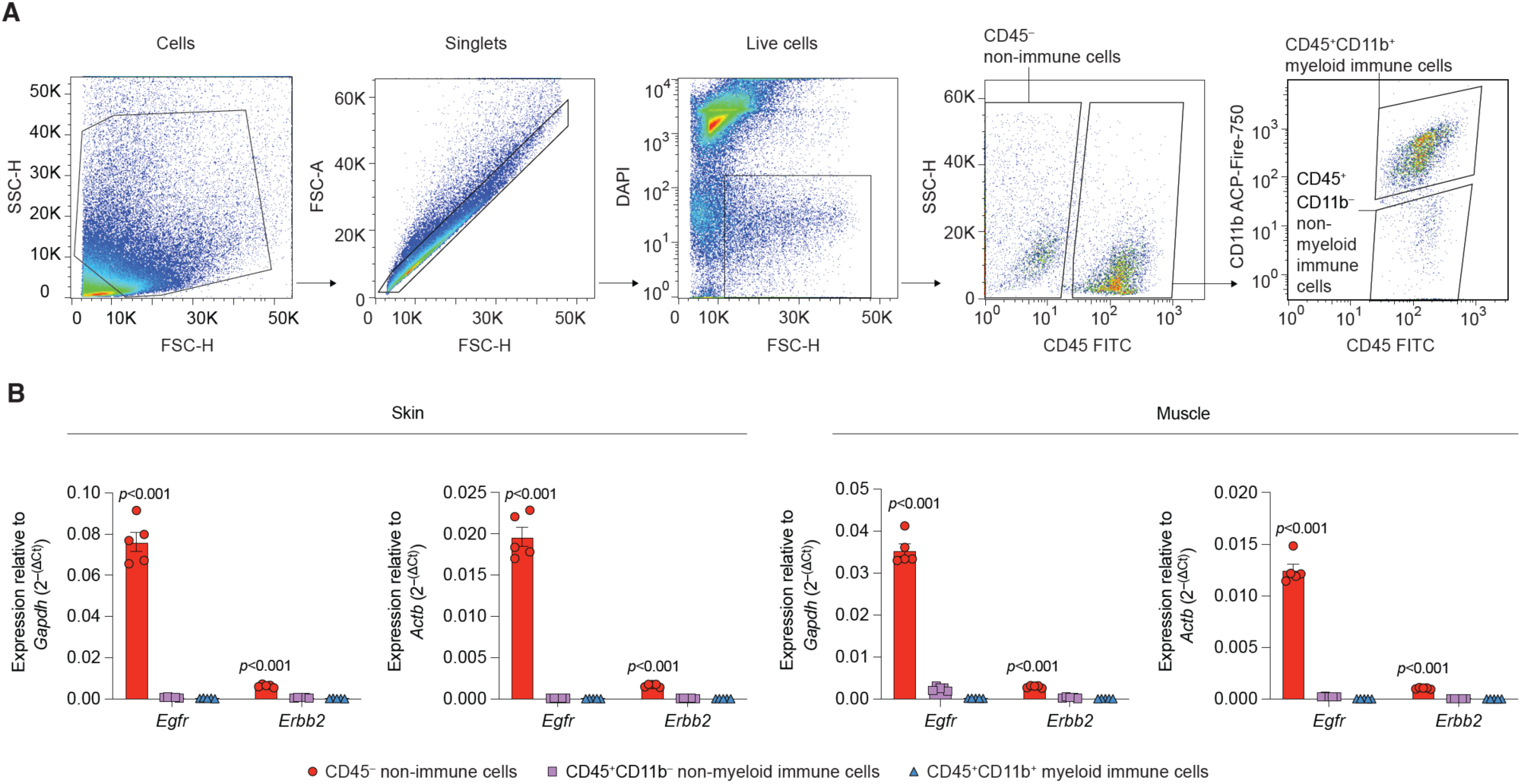
*Egfr* and *Erbb2* expression in immune cells from injured tissues is minimal. (**A**) Sequential gating strategy to sort non-immune cells (CD45^−^), non-myeloid immune cells (CD45^+^ CD11b^−^), and myeloid immune cells (CD45^+^ CD11b^+^) from injured tissues 4 days post-injury. Cell debris, doublets, and dead cells were excluded prior to gating on the target populations. (**B**) Relative expression of *Egfr* and *Erbb2* in the sorted populations was assessed by qPCR relative to the housekeeping genes *Gapdh* and *Actb*. Data are presented as mean ± SEM. Statistical significance was determined using one-way ANOVA with Bonferroni post hoc tests. *P* values are indicated.

**Supplementary Figure 7.**
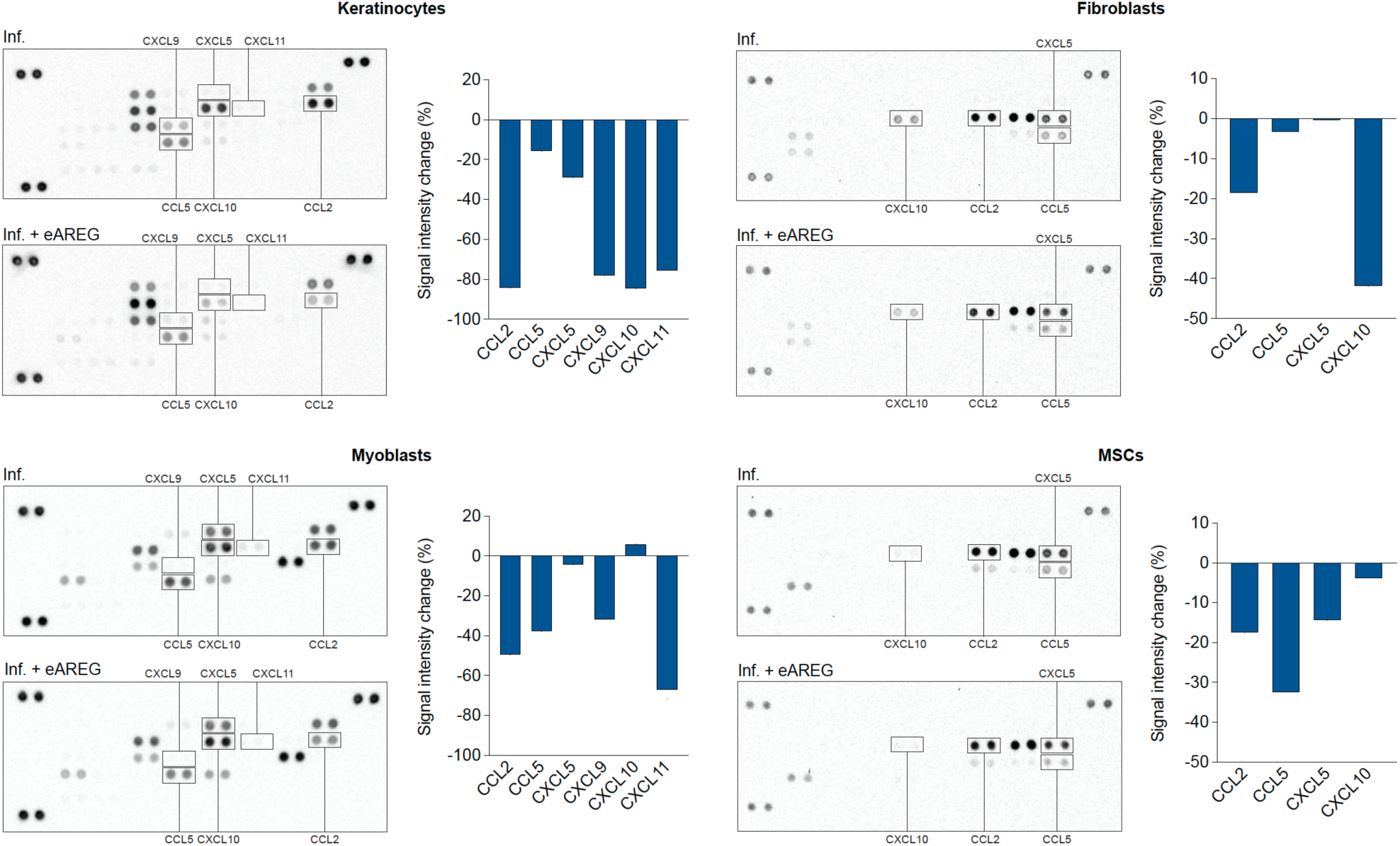
eAREG suppresses chemokine production by tissue cells. Keratinocytes, fibroblasts, myoblasts MSCs were treated with an inflammatory cytokine cocktail comprised of TNF-α, IL-1β and IFN-γ only (“Inf.”), or in the presence of eAREG (“Inf. + eAREG”). Supernatants were collected after 24 hours and chemokine levels were analyzed using membrane antibody arrays. Densitometry analysis was performed to compare levels of detected chemokines between conditions. Membrane images are displayed on the left for each cell type, with quantification of key selected chemokines shown on the right. *n* = 2 technical replicates.

**Supplementary Figure 8.**
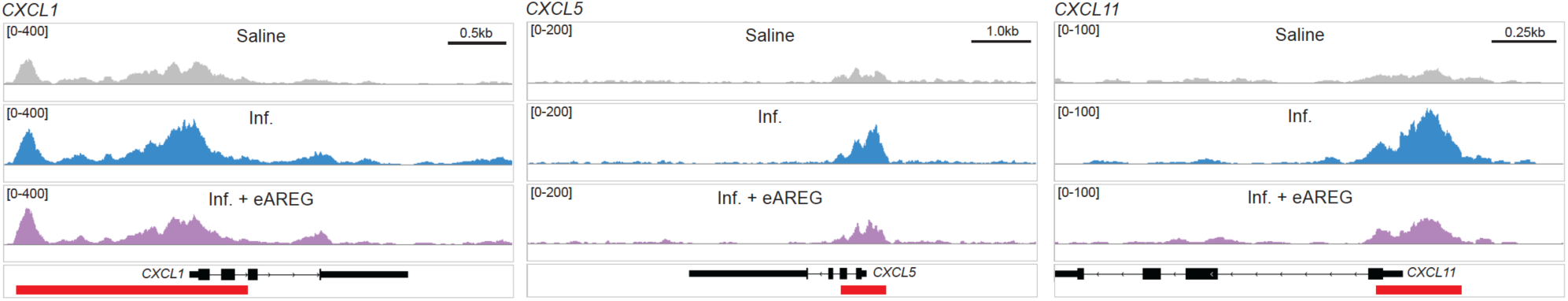
eAREG limits chemokine expression in tissue cells by decreasing chromatin accessibility. Keratinocytes were treated with saline control; an inflammatory cytokine cocktail comprised of TNF-α, IL-1β and IFN-γ only (Inf.); or inflammatory cytokine cocktail and eAREG (Inf. + eAREG). Cells were collected after 24 hours and processed for analysis by ATAC-seq. Data were processed using the *nf-core/atacseq* pipeline and regulatory regions (i.e. promoters/enhancers) were annotated using HOMER (as part of the pipeline) or identified within the GeneHancer database. Merged ATAC-seq read tracks are displayed for each treatment condition at the promoter/enhancer loci for *CXCL1*, *CXCL5*, and *CXCL11*. The schematic for each gene is displayed in black below the tracks, and the quantified peaks/regulatory regions are indicated by the red bars. Scale bars are indicated in kilobases (kb).

